# Actin disassembly triggers CNS myelin compaction and wrapping

**DOI:** 10.64898/2026.05.14.722739

**Authors:** Husniye Kantarci, Kathryn Wu, Nicholas Ambiel, Eduardo Chaparro Barriera, Madeline H. Cooper, Alexandra Münch, Yari M. Sigal, Miguel A. Garcia, Manasi Iyer, Danielle M. Jorgens, J. Bradley Zuchero

## Abstract

Compact myelin enables rapid and precise impulse conduction in the vertebrate nervous system. During CNS development, oligodendrocytes wrap spirally around axons while compacting membranes, yet how cytoskeletal remodeling is coupled to these events remains unclear. Actin disassembly is required for wrapping, but whether wrapping is driven by iterative actin-based protrusion or follows a transition to actin-independent mechanisms is unresolved. Here we integrate tract-resolved developmental profiling, live-cell compaction mapping, and in vivo genetic perturbation to define how actin disassembly is coupled to myelin wrapping. We find that actin filaments undergo pronounced, sustained disassembly before active wrapping begins, with little evidence for persistent actin within sheaths during wrapping. We develop a live-cell assay that maps membrane compaction in cultured oligodendrocytes and show that compaction zones are depleted of actin filaments and expand when actin disassembly is promoted. Consistent with this model, oligodendrocyte-specific actin disassembly in vivo accelerates the appearance of thicker myelin early in development. Together, our results support a model in which actin disassembly promotes myelin wrapping by enabling membrane compaction, and provide a platform to dissect how compaction is regulated in development and disease.

## INTRODUCTION

Myelin enables rapid and precise signal propagation throughout the vertebrate nervous system and is critical for normal development, learning, and neuronal survival^1–3^. During CNS myelin formation, oligodendrocytes first ensheath axons with several circumferential turns of cytoplasm-rich processes^4–7^. They subsequently generate the remaining bulk of myelin layers through spiral wrapping while concomitantly compacting their membranes to form mature, compact myelin^8^. This second phase of wrapping and compaction establishes myelin’s defining role as an electrical insulator^9^ by increasing the resistance and decreasing the capacitance across the axonal membrane^10^. This multilamellar wrapping is a universal mechanism across jawed vertebrates to facilitate rapid conduction through saltatory propagation^11^. However, the molecular mechanisms that orchestrate compaction and wrapping remain inadequately understood.

Myelination requires extensive cytoskeletal remodeling within oligodendrocyte processes^12,13^. While ensheathment relies on branched actin filament assembly at the leading edge of the oligodendrocyte process^14^, wrapping is surprisingly characterized by rapid and widespread actin disassembly^14,15^. Oligodendrocytes upregulate actin disassembly factors such as ADF, cofilin-1, and gelsolin as they mature, and compact myelin is largely devoid of actin filaments^14,15^. Genetic deletion of the actin disassembly factors gelsolin^14^ or ADF and cofilin1^15^ in mice reduces the number of myelin wraps, whereas pharmacological promotion of actin disassembly with latrunculin increases wraps^14^. Together, these results indicate that actin disassembly occurs during myelin wrapping and is both necessary and sufficient for this process. This raises a central mechanistic question: how does actin disassembly promote myelin wrapping^16^?

Current models of myelin wrapping fall into two competing categories: actin-dependent and actin-independent mechanisms^13–16^. The main actin-dependent hypothesis, which we refer to here as the “actin cycling model,” posits that repeated cycles of actin assembly and disassembly at the leading edge incrementally advance the oligodendrocyte process around the axon (potentially in a “ratcheting” manner)^15^. Because myelin wrapping proceeds normally in mice lacking the Arp2/3 complex^14^—thereby ruling out canonical lamellipodial protrusion—such cycling would require a non-canonical mode of actin-based protrusion. In contrast, the primary actin-independent hypothesis, which we call the “trigger model,” proposes that actin disassembly initiates subsequent steps in wrapping without requiring sustained actin assembly at the leading edge^14^. The central distinction between these models is whether ongoing actin assembly is required during wrapping. Resolving this question requires defining the temporal relationship between actin filaments (F-actin) and the onset and progression of wrapping: persistent F-actin within sheaths during wrapping would support the actin cycling model, whereas widespread loss of F-actin prior to or during wrapping would favor a trigger mechanism.

If, as suggested by the trigger model, actin disassembly does not advance wrapping through iterative actin cycling, how does it contribute to myelin wrapping? We hypothesize that actin disassembly enables myelin compaction, a process tightly coupled to wrapping^6,7^ and likely necessary for continued spiral growth^17,18^. During compaction, myelin basic protein (MBP) mediates intracellular adhesion between oligodendrocyte membranes and progressively “zippers” the sheath from the outer layers toward the inner tongue^19,20^. The vast majority of cytoplasm and proteins are displaced from compact myelin, which retains only an ∼3 nm intracellular compartment largely populated by MBP^7,21^. Because actin filaments are 7 nm in diameter, even sparse F-actin would be expected to sterically hinder intracellular membrane apposition, suggesting that actin disassembly could be permissive for compaction^14,15^. However, whether actin disassembly directly promotes compaction has not been experimentally tested.

Here, we show that actin disassembly promotes myelination by facilitating membrane compaction. By defining the developmental timing of actin loss relative to myelin ultrastructure, we find that actin filaments disappear from nascent sheaths as wrapping proceeds, arguing against a requirement for sustained actin assembly during active wrapping. We further introduce a live-cell assay to map MBP-associated compaction in cultured oligodendrocytes and demonstrate that experimentally promoting actin disassembly increases compaction in culture. Finally, we use oligodendrocyte-restricted expression of DeAct genetic tools to show that actin disassembly accelerates developmental myelin wrapping in vivo.

## RESULTS

### Myelin wrapping in the mouse CNS is protracted and slow

To test whether myelin wrapping advances through iterative actin-based protrusion (the actin cycling model) or instead follows a transition to actin-independent mechanisms (the trigger model) (Supplementary Fig. 1), we first sought to establish when myelin sheaths are predominantly in the <u>ensheathment</u> versus the <u>wrapping</u> phase. Previous studies in the developing mouse spinal cord reported bright F-actin foci in nascent MBP+ myelin profiles at early postnatal stages^7,14,15^, but it has remained unclear whether these actin structures are present during ensheathment, subsequent spiral wrapping, or both. Interpreting actin localization, therefore, requires a tract-resolved developmental time course that defines when ensheathment transitions to wrapping.

Because the time course of myelin formation varies by CNS region, even within the spinal cord^6,22,23^, we focused on a single, anatomically defined white matter tract: the thoracic gracile fasciculus (Fig. 1a). This dorsal column tract contains densely packed, largely parallel ascending sensory axons^22^, enabling quantitative analysis of hundreds of myelin cross-sections per animal across development.

**Figure 1.**
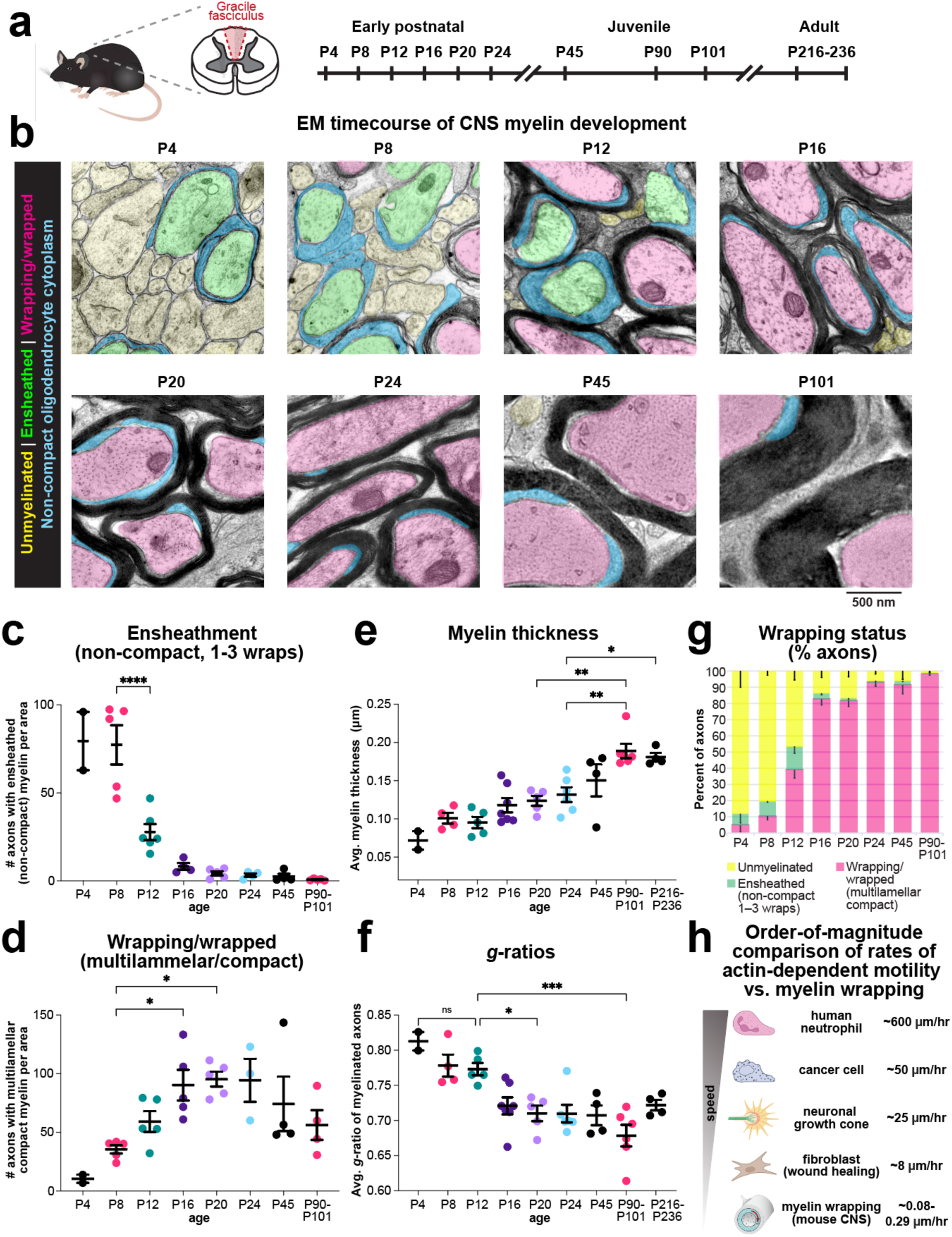
Myelin wrapping in the mouse spinal cord is developmentally staged and prolonged. **a,** Schematic of the mouse thoracic spinal cord showing the gracile fasciculus region analyzed (left) and sampling timeline (right). Axis breaks indicate omitted intervals between early postnatal, juvenile, and adult ages. **b,** Representative transmission electron micrographs from an EM time course of gracile fasciculus development. Axons were classified as unmyelinated (yellow), ensheathed (green; 1–3 non-compact wraps), or wrapping/wrapped (magenta; multilamellar compact, major dense line-positive multilayered myelin). Non-compact oligodendrocyte cytoplasm is indicated (cyan). All color is false-coloring; raw micrographs are shown in Supplementary Fig. 2. Scale bar, 500 nm. **c,** Quantification of ensheathed axons (non-compact, 1–3 wraps) per analyzed area across development (EM), normalized to a region of 194.1882 µm^2^. P4 (N = 2 mice), P8 (N = 5 mice), P12 (N = 6 mice), P16 (N = 4 mice), P20 (N = 5 mice), P24 (N = 4 mice), P45 (N = 4 mice), and P90-101 (N = 5 mice). **d,** Quantification of axons classified as wrapping/wrapped (multilamellar compact myelin) per analyzed area across development (EM), normalized to a region of 194.1882 µm^2^. P4 (N = 2 mice), P8 (N = 5 mice), P12 (N = 5 mice), P16 (N = 5 mice), P20 (N = 5 mice), P24 (N = 3 mice), P45 (N = 4 mice), and P90-101 (N = 4 mice). **e,** Average myelin thickness across development (EM). P4 (N = 2 mice), P8 (N = 5 mice), P12 (N = 5 mice), P16 (N = 7 mice), P20 (N = 5 mice), P24 (N = 6 mice), P45 (N = 4 mice), P90-101 (N = 6 mice), and P216-236 (N=4 mice). **f,** Average *g*-ratio across development (EM). p4 (N = 2 mice), P8 (N = 5 mice), P12 (N = 5 mice), P16 (N = 7 mice), P20 (N = 5 mice), P24 (N = 6 mice), P45 (N = 4 mice), P90-101 (N = 6 mice), and P216-236 (N=4 mice). **g,** Wrapping status across development shown as the percent of axons per animal classified as unmyelinated, ensheathed, or wrapping/wrapped (EM; percentages computed per animal and then averaged). P4 (N = 2 mice), P8 (N = 4 mice), P12 (N = 5 mice), P16 (N = 7 mice), P20 (N = 6 mice), P24 (N = 5 mice), P45 (N = 4 mice), and P90 (N = 5 mice). **h,** Order-of-magnitude comparison of reported net advance rates for actin-dependent cell motility and estimated myelin wrapping/thickening rates in the mouse CNS (see Methods for calculation and references). Data in c–f are shown as mean ± SEM with individual animals plotted as points. Statistics: ordinary one-way ANOVA with Tukey test.

We generated an ultrastructurally preserved electron microscopy time course of gracile fasciculus development from early postnatal stages (P4) through adulthood (P216–236) (Fig. 1a,b, and Supplementary Fig. 2). We classified axons as unmyelinated, ensheathed (1–3 loose, non-compact wraps^6,7,22,24^), or wrapping/wrapped (multilamellar compact myelin; major dense line-positive^7,14,24^) (Fig. 1b). Ensheathed profiles were most frequent early (P4–P8) and then declined sharply (Fig. 1c), indicating that ensheathment is a brief intermediate step, as previously inferred^6,25,26^. In contrast, the number of axons bearing multilamellar compact myelin (“wrapping/wrapped”) increased rapidly during the second and third postnatal weeks (Fig. 1d), and by P16–P24 the majority of axons were classified as wrapping/wrapped (multilamellar compact) by this ultrastructural criterion (Fig. 1b,g).

Despite the early appearance of multilamellar compact myelin, myelin growth continued for months. Average myelin thickness increased progressively from early postnatal stages into adulthood (Fig. 1e), accompanied by a corresponding decrease in *g*-ratio (Fig. 1f). Restricting analysis to the thickest 10% of sheaths to minimize bias from newly-forming sheaths, we observed that estimated myelin spiral length increased gradually across development in a pattern well described by a logarithmic function (R² = 0.91, Supplementary Fig. 2c-f).

Differentiating the fitted curve revealed that total spiral-length growth rate declined from a maximum of 0.49 μm/hr in early development to a mean of 0.15 μm/hr across the measurement period. When we partitioned spiral length increases into contributions from new wraps and radial expansion, new-wrap addition accounted for about half of the spiral length increase, declining from a maximum rate of 0.29 μm/hr to a mean of 0.081 μm/hr (R² = 0.90; Supplementary Fig. 2g; see Methods). These sustained but decelerating rates over >200 days demonstrate that myelin wrapping in the developing mouse spinal cord is a prolonged and comparatively slow process relative to canonical actin-driven motility (Fig. 1h). This tract-resolved EM time course provides the developmental staging used below to relate actin localization to ensheathment versus multilamellar myelin wrapping.

### Actin filament disassembly in nascent myelin sheaths precedes myelin wrapping

With the developmental timing of ensheathment versus multilamellar compact sheath formation in the thoracic gracile fasciculus established (Fig. 1), we next asked when and where actin filaments are present within nascent myelin sheaths. In a parallel set of animals collected at the same postnatal ages as our EM time course, we stained tissue with phalloidin to label F-actin^14^ and immunostained for myelin basic protein (MBP), which is expressed in newly differentiated oligodendrocytes^27^ and accumulates in nascent sheaths^7,14^. At early ages we frequently observed discrete F-actin foci colocalizing within MBP+ sheath profiles (Fig. 2a; see Methods). Quantification revealed that ∼50% of MBP+ sheaths contained F-actin foci at P4 and ∼36% at P8 (Fig. 2b), consistent with prior observations in the developing spinal cord^7,14,15^. In contrast, by P12, actin foci were rarely detected within MBP+ sheaths and remained low thereafter (Fig. 2b).

**Figure 2.**
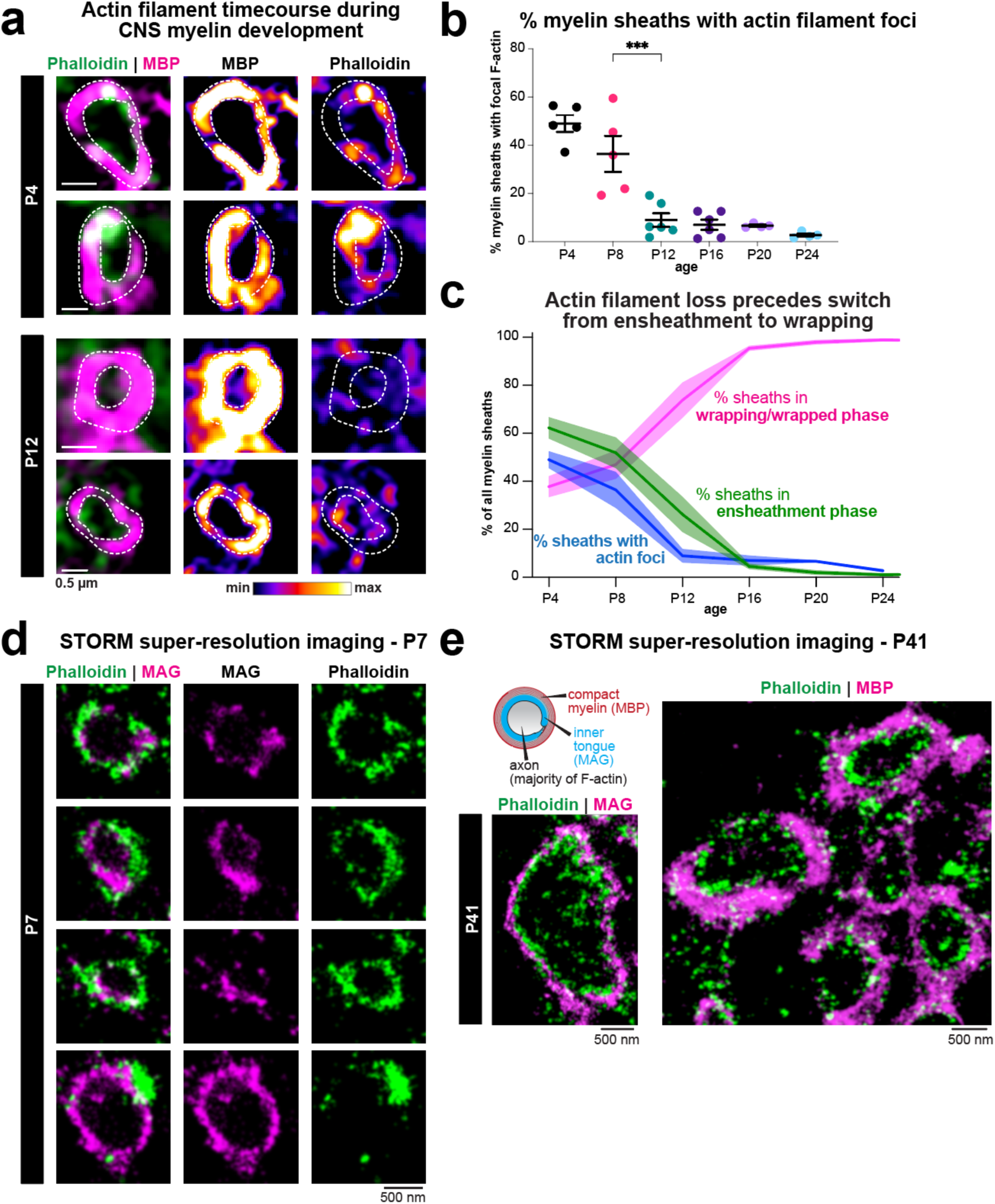
Actin filament disassembly in nascent myelin sheaths precedes myelin wrapping. **a**, Representative images of nascent myelin sheaths in the thoracic gracile fasciculus across postnatal development stained with phalloidin (F-actin) and immunolabeled for MBP. Dashed outlines indicate MBP+ sheath profiles scored for actin filament foci. Scale bar, 0.5 µm. **b**, Quantification of the percentage of MBP+ myelin sheath profiles containing discrete actin filament foci across development. Each dot represents the mean per animal; bars show mean ± SEM. Statistics: ordinary one-way ANOVA with Tukey test. P4 (N = 5 mice), P8 (N = 5 mice), P12 (N = 6 mice), P16 (N = 6 mice), P20 (N = 4 mice), and P24 (N = 4 mice). **c**, Alignment of developmental changes in actin foci with sheath ultrastructural stage (based on EM data from Figure 1). Lines show the percentage of sheaths with actin foci (blue) and the percentage of sheaths classified as being in the ensheathment phase (green) or wrapping/wrapped phase (magenta; multilamellar compact) across age (per-animal percentages). Thick lines indicate the mean across animals and shaded regions indicate SEM. **d**, STORM super-resolution imaging at P7 showing phalloidin-labeled F-actin relative to the inner tongue marker MAG in nascent sheaths. Scale bar, 500 nm. **e**, STORM super-resolution imaging at P41 showing phalloidin-labeled F-actin relative to MAG (left) and MBP (right). Cartoon indicates relative positions of axon (majority of F-actin), inner tongue (MAG), and compact myelin (MBP). Scale bars, 500 nm.

To relate the disappearance of actin foci to sheath ultrastructural stage, we aligned the actin-foci time course with our EM-based classification of axons as ensheathed (1–3 non-compact wraps) versus wrapping/wrapped (multilamellar compact). Strikingly, the decline in sheaths with actin foci preceded the population transition from the ensheathment phase to the wrapping phase (multilamellar compact) (Fig. 2c), placing F-actin loss upstream of widespread multilamellar myelin growth in this tract.

As a complementary measurement, we quantified phalloidin signal within MBP ring ROIs across development and found that background-subtracted F-actin signal within these MBP-defined sheath profiles peaked early and declined with age (Supplementary Fig. 3b) as we previously reported^14^. This analysis was robust to the choice of phalloidin intensity threshold used to define actin-positive signal (Supplementary Fig. 3c; see Methods).

Because diffraction-limited imaging cannot reliably distinguish oligodendrocyte F-actin in the inner tongue from the abundant submembranous actin present in axons^28,29^, we next used stochastic optical reconstruction microscopy (STORM)^28,30^ to localize F-actin relative to the inner tongue marker MAG. At P7, when ensheathment is still common, STORM revealed frequent F-actin enrichment adjacent to MAG+ sheath structures, consistent with actin accumulation at the leading edge during ensheathment (Fig. 2d). By P41, when ensheathment is rare, the majority of F-actin signal formed a narrow ring internal to, and largely non-overlapping with, MAG and MBP, consistent with predominantly axonal F-actin at this stage (Fig. 2e).

Together, these data support a “trigger model” in which actin filament disassembly in nascent sheaths precedes the transition from ensheathment to myelin wrapping in vivo.

### Membrane compaction mapping enables live measurement of nanoscale thinning in cultured oligodendrocytes

Having established that actin disassembly precedes wrapping in vivo, we next sought to directly test how actin filaments regulate membrane compaction—a process we hypothesize is enabled by actin loss. Cultured oligodendrocytes elaborate large, flattened membrane sheets that recapitulate several hallmark features of myelinogenesis, including induction of myelin genes such as MBP^31^, extensive cytoskeletal remodeling^14,15^, and the formation of cytoplasmic channels^31,32^. Prior work showed that MBP can self-associate^21^ and generate protein-excluded membrane domains in cultured oligodendrocytes^31^, suggesting that aspects of compaction can be modeled in culture (Supplementary Fig. 4a). However, whether these domains achieve the nanometer-scale thinning^33^ expected for close cytoplasmic apposition under physiological conditions has remained unclear, possibly because prior (atomic force microscopy, AFM) measurements were performed on permeabilized and immunostained cells^31^.

To address this, we purified primary oligodendrocyte precursor cells (OPCs)^14,34^ and expressed a membrane-anchored fluorescent reporter (EGFP-CAAX) under the MBP promoter^35–37^, together with labeling of the plasma membrane using MemGlow, a lipophilic and fluorogenic membrane dye that labels the plasma membrane independent of protein content^38,39^. As they differentiated and matured in culture, oligodendrocytes developed discrete membrane domains that were depleted of EGFP-CAAX^31^ yet remained MemGlow-positive, indicating continuous membrane regions rather than gaps or holes (Fig. 3a). We refer to these EGFP-CAAX-excluded, MemGlow-positive cellular regions as compaction zones, and to doubly labeled cells as compaction-mapped oligodendrocytes (Fig. 3a–c). We next used correlative fluorescence imaging and bio-atomic force microscopy (bio-AFM) on oligodendrocytes to directly measure membrane height in the same regions classified by fluorescence (Fig. 3d,e; Supplementary Fig. 4b). EGFP-CAAX-excluded domains were markedly thinner than surrounding membrane and fell within a ∼12–16 nm height range, consistent with nanoscale membrane apposition (Fig. 3e,f). Across line profiles, compact regions were associated with reduced EGFP-CAAX signal compared to adjacent non-compact regions (Fig. 3f and Supplementary Fig. 4c). Consistent with the interpretation that these zones reflect MBP-associated^40^ compaction-like membrane states^31^, MBP immunostaining was enriched in EGFP-CAAX-excluded domains (Supplementary Fig. 4a). Measured height values also closely match thickness estimates for compact myelin measured by electron microscopy in vivo^6,17^.

**Figure 3.**
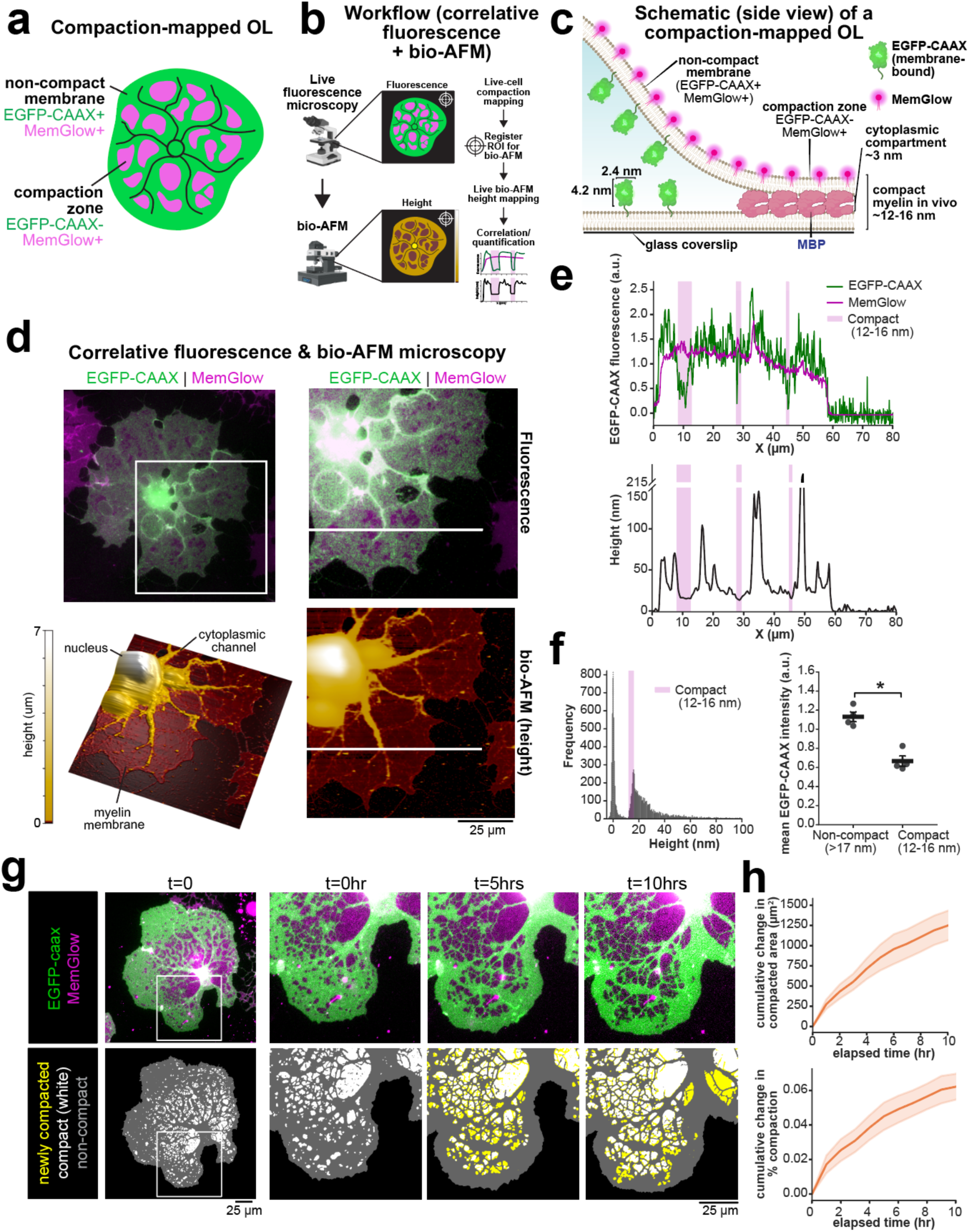
Membrane compaction mapping enables live measurement of nanoscale membrane thinning in cultured oligodendrocytes. **a**, Schematic defining compaction-mapped oligodendrocytes (OLs). Non-compact membrane regions are EGFP-CAAX+ and MemGlow+, whereas compaction zones are EGFP-CAAX− and MemGlow+. **b**, Workflow for correlative live fluorescence compaction mapping and bio-AFM height measurements in the same oligodendrocyte membrane region. |**c**, Schematic (side view) of a compaction-mapped OL highlighting EGFP-CAAX exclusion from compacted membrane and expected nanoscale thickness of compaction zones, based on known in vivo dimensions. **d**, Correlative fluorescence imaging and bio-AFM of a 3-day-differentiated OL expressing EGFP-CAAX and stained with MemGlow. The boxed region is magnified (right). Corresponding bio-AFM height maps are shown. Scale bar, 25 µm. **e**, Line scans from the region in d used to extract EGFP-CAAX and MemGlow fluorescence intensity (top) and membrane height profiles (bottom). Shading denotes compact-height regions (12–16 nm). **f**, Distribution of membrane height values measured by bio-AFM (left) and quantification of EGFP-CAAX fluorescence in non-compact (>17 nm) versus compact (12–16 nm) regions (right). Mean ± SEM; N = 4 independent biological replicates; 3 line profiles from 1–2 cells per replicate. Paired two-tailed t-test: t(3) = 5.24, p = 0.014 (exact p = 0.0135), mean difference = 0.46 (95 % CI [0.18, 0.75]). **g**, Time-lapse imaging of a representative 3-day-differentiated compaction-mapped OL. Bottom row shows segmentation of compact (white) and non-compact (gray) membrane regions by EGFP-CAAX exclusion; yellow indicates newly compacted membrane relative to the previous frame. Scale bar, 25 µm. **h**, Cumulative change in compacted area per cell (top) and cumulative change in % compaction (bottom) over time in 3-day-differentiated compaction-mapped OLs. Thick line indicates mean and shaded region indicates SEM (N = 3 biological replicates).

Because both EGFP-CAAX and MemGlow can be imaged in live cells, we next asked whether compaction zones are dynamic. Time-lapse imaging of 3-day-differentiated compaction-mapped oligodendrocytes revealed that compaction zones formed, expanded, and remodeled over hours, enabling quantification of the cumulative change in compacted membrane area and percent compaction over time (Fig. 3g,h; Supplementary Video 1). Under gentle illumination conditions, we could routinely live-image oligodendrocytes for 10+ hours with 1 frame/hr frame rates. On average, oligodendrocytes compacted their membranes at a rate of 132 µm²/hr, increasing the proportion of total membrane area compacted by 0.6% per hour.

Compaction was surprisingly dynamic, and individual compaction zones occasionally decompacted even as cells underwent net compaction overall (Supplementary Video 2). Overall, compaction mapping provides a live-cell readout of membrane domains that exhibit nanoscale thinning and dynamic growth, creating a platform to test how cytoskeletal remodeling regulates compaction.

### Actin disassembly promotes expansion and fusion of compaction zones in cultured oligodendrocytes

We next used compaction mapping to ask how actin organization relates to membrane compaction and whether actin disassembly is sufficient to promote it. In 7-day-differentiated compaction-mapped oligodendrocytes, phalloidin staining revealed that F-actin was reduced within compaction zones relative to adjacent non-compact membrane (Fig. 4a,b). Live imaging with the genetically-encoded actin reporter Lifeact-mRuby3^41,42^ further showed persistently lower actin signal in compact regions over time (Fig. 4c; Supplementary Fig. 5a-b; Supplementary Video 3), supporting the conclusion that compacted membrane domains are F-actin-depleted in both fixed and live assays. To determine whether actin disassembly or clearance preceded compaction, we analyzed Lifeact intensity in membrane regions that subsequently became compacted. Compared to neighboring regions that never compacted, regions destined to compact showed a trend towards lower Lifeact intensity, which decreased sharply at compaction onset (Supplementary Fig. 5c). As our imaging interval was 1 hour, our data suggest that actin disassembly/clearance occurs within the hour preceding compaction or at the time of compaction.

**Figure 4.**
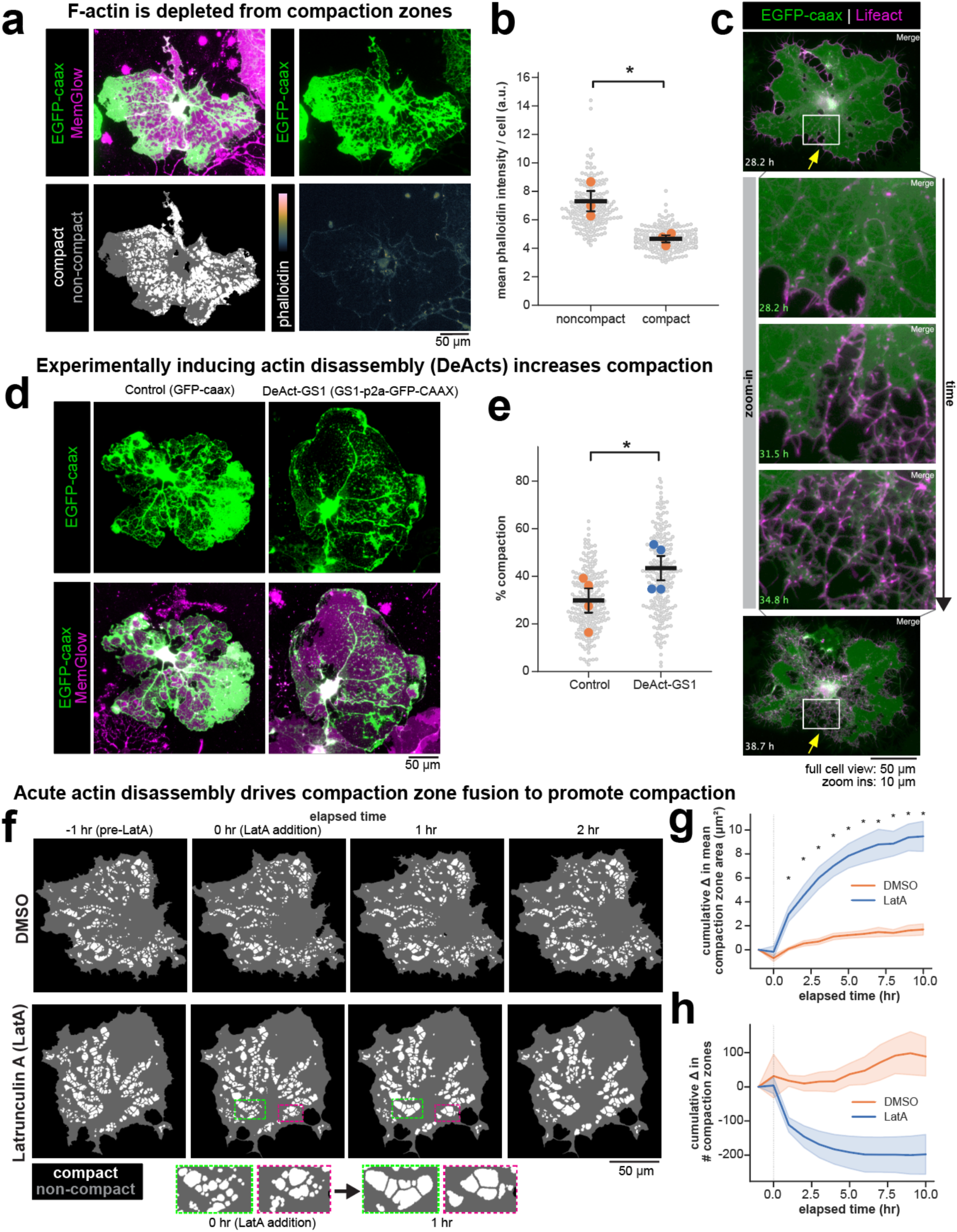
Actin disassembly promotes expansion and fusion of compaction zones. **a**, F-actin distribution in 7-day-differentiated compaction-mapped OLs stained with phalloidin. EGFP-CAAX/MemGlow images, segmentation of compact (white) and non-compact (gray) membrane regions, and phalloidin intensity (BatlowK colormap) are shown. Scale bar, 50 µm. **b**, Quantification of phalloidin intensity in compact versus non-compact membrane regions. Phalloidin signal was reduced in compact regions. Mean ± SEM; N = 3 independent biological replicates. Paired two-tailed t-test: t(2) = 5.35, p = 0.033, mean difference = −2.64 (95% CI [−4.77, −0.52]). **c**, Live imaging of a compaction-mapped OL expressing Lifeact-mRuby3 (Lifeact) showing low Lifeact signal in compact zones and a decrease in Lifeact intensity coincident with compaction onset in the boxed region. Scale bars: full cell view, 50 µm; zoom-in, 10 µm. **d**, Representative images of 7-day-differentiated compaction-mapped OLs expressing control (EGFP-CAAX) or DeAct-GS1 (GS1-P2A-EGFP-CAAX). Scale bar, 50 µm. **e**, Quantification of percent compaction per cell in control versus DeAct-GS1–expressing OLs. Mean ± SEM; N = 4 independent biological replicates. Paired two-tailed t-test: t(3) = 5.36, p = 0.013, mean difference = 0.14 (95% CI [0.06, 0.22]). **f**, Compaction masks from live imaging before and after latrunculin A (LatA) addition (0 h) compared with DMSO control, showing compaction zone remodeling and fusion. Scale bar, 50 µm. Insets highlight representative fusion events. See also Supplementary Video 4. **g**, Cumulative change in mean compaction zone area over time after LatA or DMSO addition. Thick line indicates mean and shaded region indicates SEM. Two-way repeated-measures ANOVA; N = 3 independent biological replicates: main effect of time, p = 1.0 × 10⁻¹³; main effect of group, p = 0.013; time × group interaction, p = 2.1 × 10⁻⁷. Per-timepoint paired two-tailed t-tests, Benjamini–Hochberg FDR-corrected: significant at t = 1–10 h (p_FDR = 0.044, 0.044, 0.044, 0.044, 0.044, 0.044, 0.044, 0.044, 0.044, 0.044); not significant at t = 0 h (p_FDR = 0.37). **h**, Cumulative change in the number of compaction zones over time after LatA or DMSO addition. Thick line indicates mean and shaded region indicates SEM. Two-way repeated-measures ANOVA; N = 3 independent biological replicates: main effect of time, p = 0.032; main effect of group, p = 0.052 (n.s.); time × group interaction, p = 5.2 × 10⁻⁵. Per-timepoint paired two-tailed t-tests, Benjamini–Hochberg FDR-corrected: no timepoints significant after correction (smallest p_FDR = 0.051 at t = 1–5 h).

To test whether F-actin loss causes compaction, we promoted actin disassembly in differentiating oligodendrocytes using DeAct-GS1 (a genetically-encoded tool to induce actin disassembly in cells^37,43^; MBP promoter drives increasing expression starting at 2d differentiation^37^). DeAct-GS1 expression from days 2–7 of differentiation reduced cellular F-actin (see Refs. ^37,43^ and Supplementary Fig. 6a) and increased the fraction of membrane area classified as compact compared with control cells that expressed only EGFP-CAAX (Fig. 4d,e). Finally, to determine how acute actin disassembly affects compaction dynamics, we treated oligodendrocytes with latrunculin A (LatA) during live imaging. LatA rapidly increased compaction zone remodeling characterized by merging/fusion of existing zones (Fig. 4f), producing an increase in the cumulative change in compaction zone area and a corresponding reduction in the number of discrete compaction zones compared with DMSO controls (Fig. 4g,h). Raw fluorescence images underlying the segmentation masks are shown in Supplementary Fig. 5d.

Together, these results identify actin disassembly as a positive regulator of membrane compaction in cultured oligodendrocytes and establish a live-imaging framework to quantify compaction dynamics and their cellular control.

### Oligodendrocyte-specific actin disassembly accelerates myelin wrapping/compaction in vivo

Actin disassembly increased MBP-associated membrane compaction in cultured oligodendrocytes (Fig. 4), prompting us to test whether promoting actin filament disassembly in oligodendrocytes is sufficient to accelerate myelin wrapping/compaction in vivo. We delivered AAVs by neonatal intrathecal injection (P0–P1) to express DeAct-GS1 under the MBP promoter (pMBP) together with a membrane-targeted EGFP reporter (DeAct-GS1-P2A-EGFP-CAAX; a “global/cytoplasmic” DeAct-GS1 that self-cleaves from membrane-bound EGFP-CAAX) and compared these animals to pMBP-EGFP-CAAX controls (Fig. 5a).

**Figure 5.**
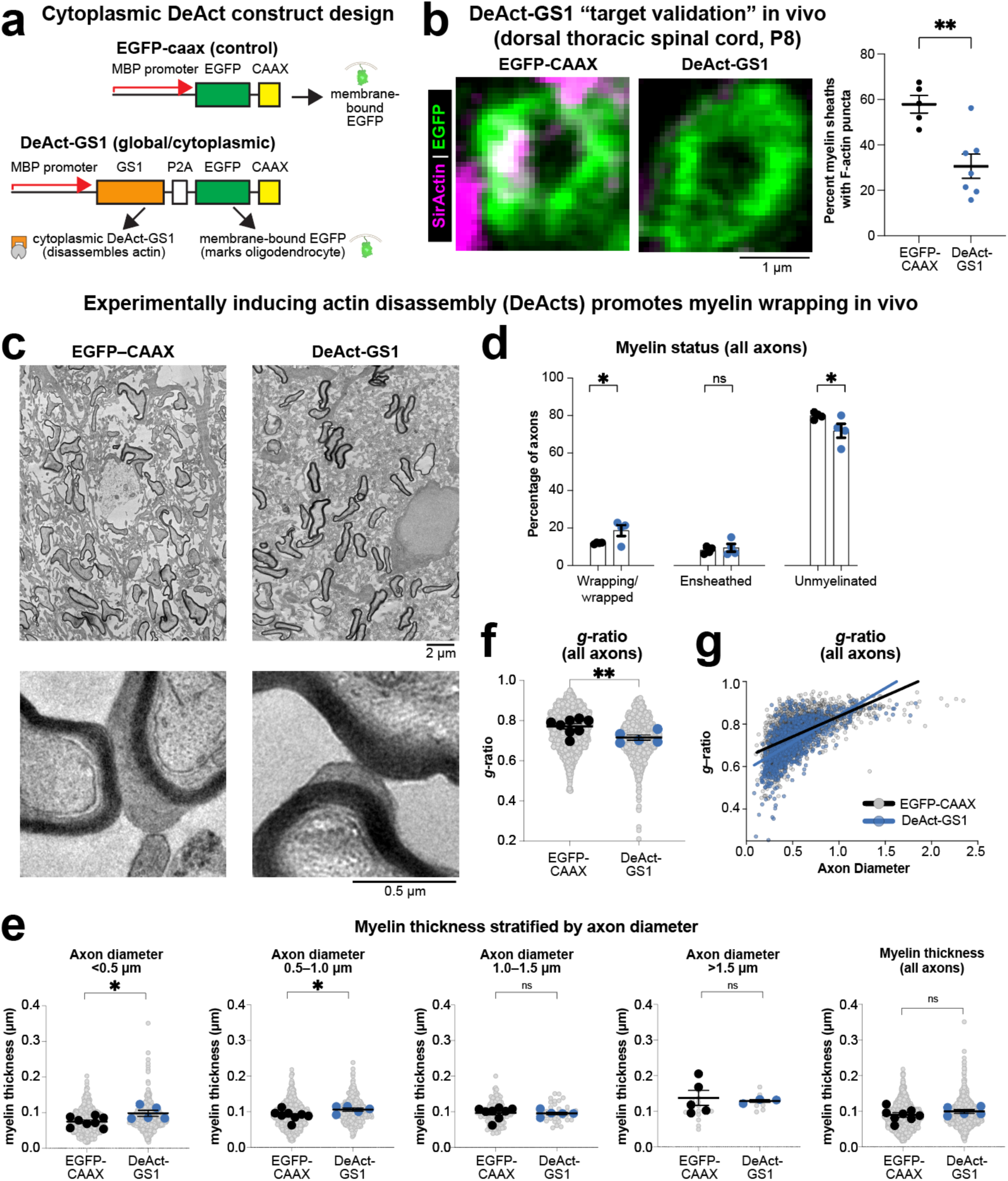
Oligodendrocyte-specific actin disassembly accelerates myelin wrapping/compaction in vivo. **a**, AAV design for oligodendrocyte-specific expression (MBP promoter). Control vector expresses membrane-targeted EGFP (EGFP-CAAX). The “global/cytoplasmic” DeAct-GS1 vector expresses gelsolin segment 1 (GS1) plus EGFP-CAAX via a “self-cleaving” P2A element (full name: pMBP-GS1-P2A-EGFP-CAAX). **b**, In vivo validation of actin disruption in myelin sheaths using SiR-actin in thick spinal cord sections at P8. Representative EGFP-labeled myelin sheath profiles from control and pMBP-GS1-P2A-EGFP-CAAX (DeAct-GS1)-injected animals are shown with corresponding quantification of the percentage of EGFP+ sheaths containing actin puncta (each dot = animal; mean ± SEM shown; n = 5–7 animals per group). Statistics: Unpaired, one-tailed t-test with Welch’s correction, **p < 0.01. Scale bar, 1 µm. **c**, Representative EM images of thoracic gracile fasciculus cross-sections at P8 from control and GS1-P2A-EGFP-CAAX (DeAct-GS1)-injected animals. Scale bars: low magnification, 2 µm; high magnification, 0.5 µm. **d**, Quantification of myelin status (wrapping/wrapped, ensheathed, or unmyelinated) at P8 in control versus DeAct-GS1 conditions. Data show mean ± SEM (n = 4 animals per group). Statistics: Unpaired, one-tailed t-test, *p < 0.05. **e**, Myelin thickness stratified by axon diameter at P8. Axons were binned by diameter (leftmost four panels) and overall thickness across all axons is shown (rightmost panel). DeAct-GS1 increased myelin thickness on small-caliber axons (diameter <1 μm) with progressively smaller effects on larger axons. SuperPlots show individual axons (small gray dots) and per-animal means (large dots); statistics were performed on per-animal means (n =5–8 animals per group, except only 3–5 animals for the >1.5 μm bin due to paucity of axons that large in some samples). Statistics: unpaired, one-tailed t-test with Welch’s correction per comparison; *p < 0.05, ns = not significant. **f**, Quantification of *g*-ratio at P8 in control versus DeAct-GS1 conditions. SuperPlots show individual axons (small gray dots) and per-animal means (large dots); statistics were performed on per-animal means (n = 5–8 animals per group). Statistics: unpaired, one-tailed t-test with Welch’s correction, **p < 0.01. **g**, *g*-ratio plotted versus axon diameter showing the relationship across axon calibers. Lines show trend for each condition. Individual axons shown as small dots; n = 5–8 animals per group.

To validate target engagement in vivo, we quantified actin puncta within EGFP-labeled myelin sheaths in thick spinal cord sections using the highly-specific^37^, fluorogenic, cell-permeable actin probe, SiR-actin^44^. At P8, ∼60% of EGFP+ sheath profiles were SiR-actin-positive in controls, whereas GS1 expression reduced this fraction to ∼30% (Fig. 5b), confirming that DeAct-GS1 disrupts sheath-associated F-actin in vivo.

We next assessed myelin ultrastructure in the thoracic gracile fasciculus at P8, a stage when many axons are transitioning from ensheathment toward active multilamellar growth (Fig. 1–2). Electron microscopy revealed thicker myelin profiles in DeAct-GS1-injected animals (Fig. 5c). DeAct-GS1 expression increased the percentage of axons classified as wrapping/wrapped (multilamellar compact myelin) with a concomitant decrease in unmyelinated axons (Fig. 5d), consistent with accelerated progression from ensheathment to active wrapping. Stratifying by axon diameter revealed that DeAct-GS1 increased myelin thickness specifically on small-caliber axons (diameter <1 μm), with progressively smaller effects on larger axons (Fig. 5e). This diameter dependence is consistent with DeAct-GS1 accelerating the onset of wrapping in axons that are just beginning to myelinate at P8, while having less effect on axons that have already accumulated multiple wraps. Correspondingly, DeAct-GS1 expression produced a pronounced shift toward lower *g*-ratios relative to controls (Fig. 5f,g). Because myelin ultrastructure was quantified across all axons in the tract (including uninfected axons), these shifts likely underestimate the cell-autonomous magnitude of the DeAct-GS1 effect. Together, these data indicate that oligodendrocyte-specific induction of actin disassembly is sufficient to accelerate myelin wrapping/compaction early in development.

### Membrane-proximal actin disassembly is sufficient to promote myelin wrapping/compaction

We next asked whether the in vivo effect of DeAct-GS1 depends on where actin disassembly is induced within the oligodendrocyte. To enrich DeAct-GS1 at the oligodendrocyte membrane, we expressed a membrane-tethered construct (GS1-EGFP-CAAX; no P2A; Fig. 6a) and assessed myelin ultrastructure at P8. Membrane-tethered GS1 increased myelin thickness and reduced *g*-ratio relative to EGFP-CAAX controls (Fig. 6b-d), with *g*-ratio shifts observed across the majority of axon diameters analyzed.

**Figure 6.**
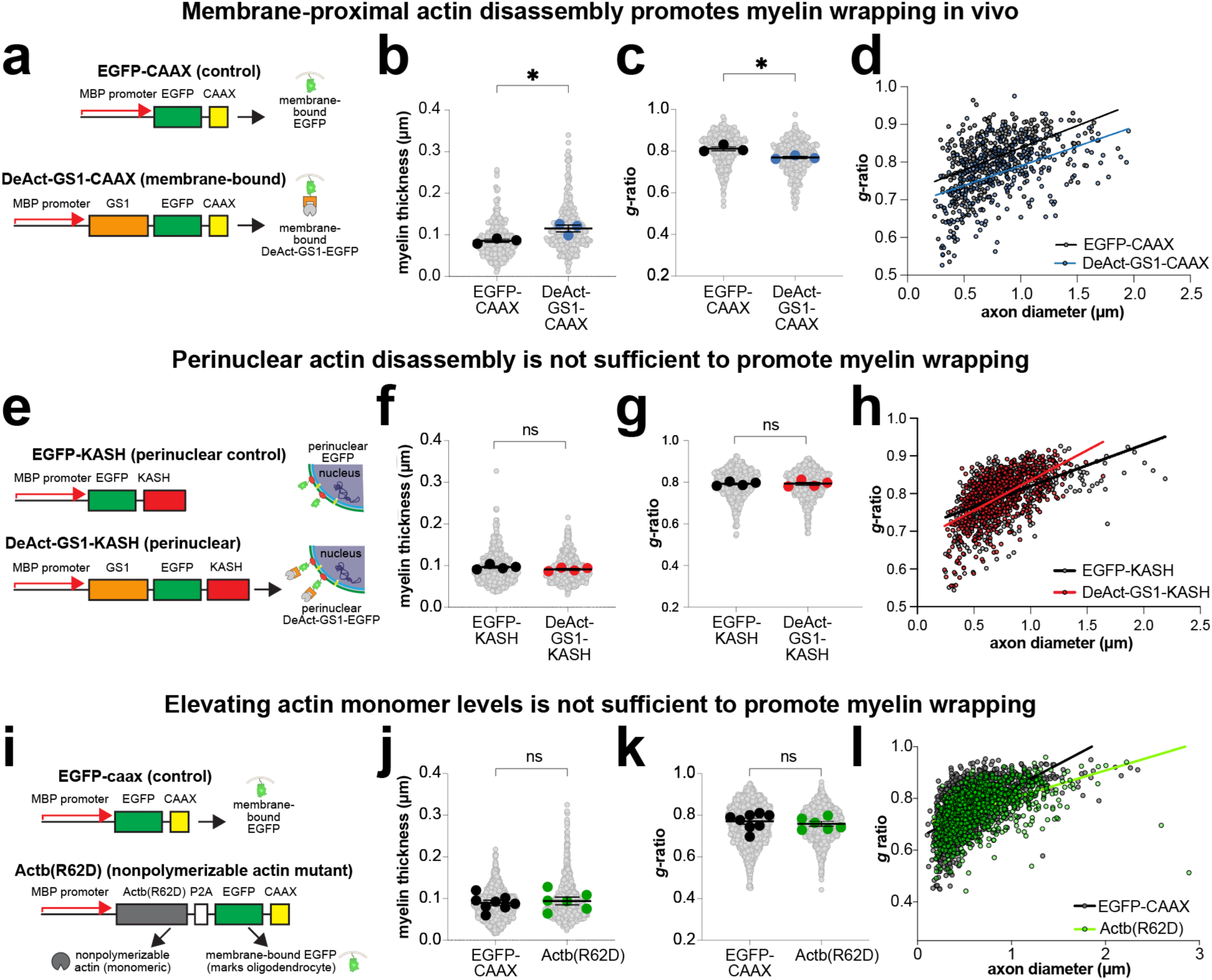
Membrane-proximal actin disassembly is sufficient to promote myelin wrapping/compaction in vivo. **a**, AAV designs used to test role of membrane-proximal F-actin in myelin wrapping. Control vector expresses membrane-targeted EGFP (EGFP-CAAX). Membrane-targeted DeAct-GS1 expresses GS1 directly fused to membrane-targeted EGFP (GS1-EGFP-CAAX; no P2A). **b-d**, Quantification of myelin thickness (b) and *g*-ratio (c) at P8 following expression of membrane-targeted DeAct-GS1-CAAX compared with EGFP-CAAX control. SuperPlots show individual axons (small gray dots) and per-animal means (large dots); statistics were performed on per-animal means (n = 3 animals per group). **d**, *g*-ratio plotted versus axon diameter with trend lines. Statistics: Unpaired, one-tailed t-test with Welch’s correction. **e**, AAV designs used to restrict actin disassembly to the oligodendrocyte cell body/outer nuclear envelope. Perinuclear-targeted DeAct expresses GS1 fused to a KASH domain (GS1-KASH) with matched EGFP-KASH control. **f-h**, Quantification of myelin thickness (f) and *g*-ratio (g) at P8 following expression of perinuclear-targeted GS1-KASH compared with EGFP-KASH control. SuperPlots show individual axons and per-animal means; statistics were performed on per-animal means (n = 4 animals per group). **h**, *g*-ratio versus axon diameter with trend lines. Statistics: Unpaired, one-tailed t-test with Welch’s correction. **i**, AAV designs to elevate monomeric actin level independent of actin disassembly. The Actb(R62D) construct expresses a nonpolymerizable mutant of beta actin plus EGFP-CAAX via a “self-cleaving” P2A element (full name: pMBP-Actb(R62D)-P2A-EGFP-CAAX). **j-l**, Quantification of myelin thickness (j) and *g*-ratio (k) at P8 following expression of Actin(R62D)-P2A-EGFP-CAAX compared with EGFP-CAAX control. SuperPlots show individual axons and per-animal means; statistics were performed on per-animal means (n = 6–8 animals per group). **l**, *g*-ratio versus axon diameter with trend lines. Statistics: Unpaired, one-tailed t-test with Welch’s correction.

To test whether actin disassembly in a non-membrane compartment is sufficient to drive wrapping/compaction, we targeted DeAct-GS1 to the perinuclear envelope^45,46^ (DeAct-GS1-KASH; Fig. 6e) and compared these animals to EGFP-KASH controls. In contrast to membrane-tethered DeAct-GS1, perinuclear DeAct-GS1-KASH had no detectable effect on myelin thickness or *g*-ratio (Fig. 6f-h), indicating that the pro-wrapping effect of actin disassembly is spatially constrained.

Finally, to distinguish actin filament disassembly from potential secondary effects of increasing actin monomer levels, we expressed a nonpolymerizable actin mutant^47^ (Actin(R62D)-P2A-EGFP-CAAX; Fig. 6i). Actin(R62D) did not alter myelin thickness or *g*-ratio relative to EGFP-CAAX controls (Fig. 6j-l), arguing that the acceleration of myelin wrapping/compaction reflects local actin filament disassembly rather than elevated monomeric actin.

Together, these in vivo perturbations show that inducing actin filament disassembly in oligodendrocytes—particularly at membrane-proximal regions—accelerates myelin wrapping/compaction during development. Along with our developmental staging and compaction-mapping experiments (Figs. 1–4), these data support a model in which actin disassembly promotes myelin wrapping by enabling membrane compaction (Fig. 7).

**Figure 7.**
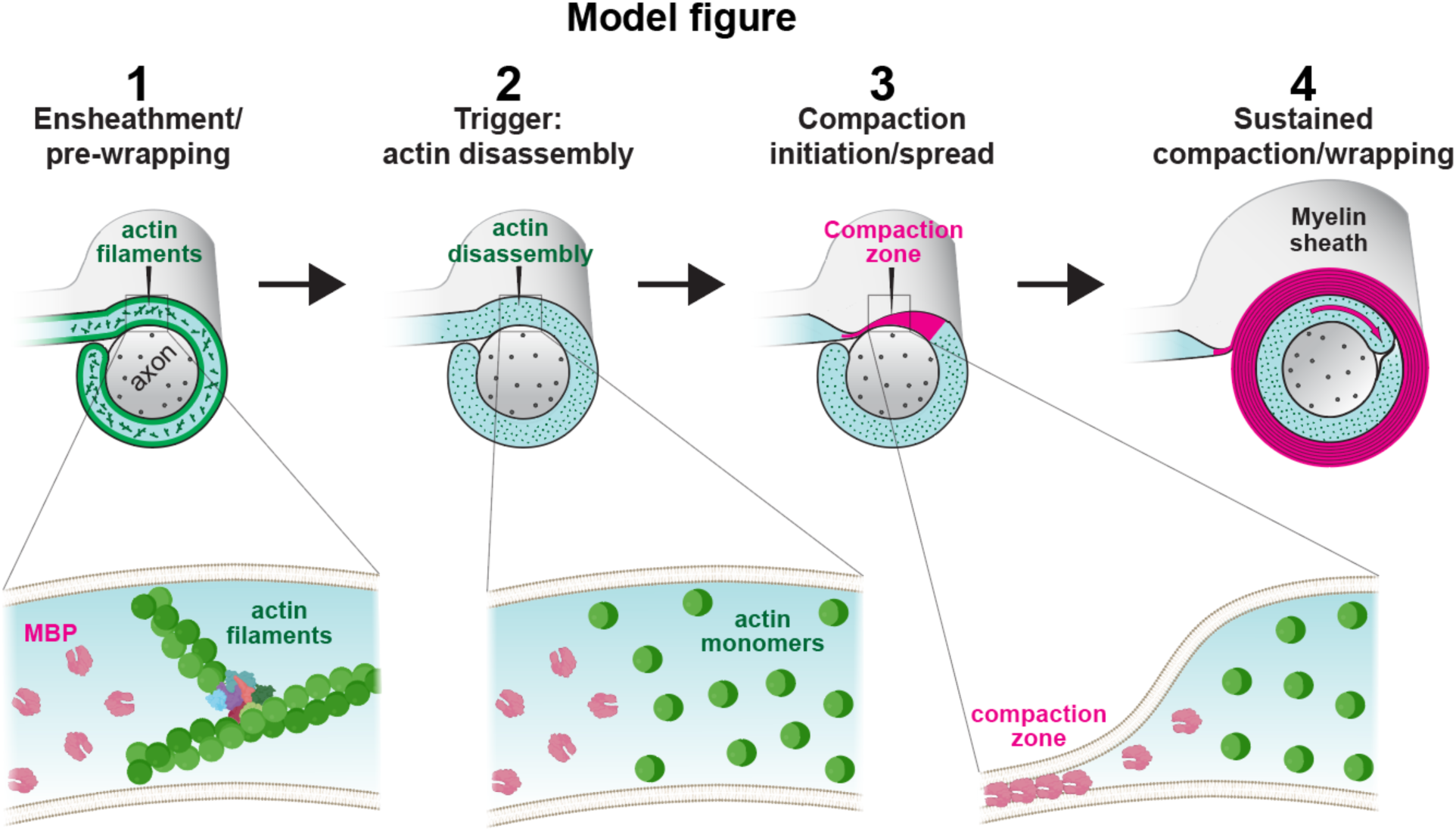
Model figure. Working model for how actin disassembly triggers myelin compaction and wrapping: (1) During ensheathment, nascent sheaths contain actin filaments. (2) A sustained actin disassembly event (“trigger”) precedes active compaction/wrapping. (3) Loss of actin filaments is permissive for membrane compaction by MBP, which forms dynamic compaction zones that are depleted of actin filaments and expand upon actin disassembly. (4) Compaction is proposed to support continued wrapping, producing thicker myelin; membrane-proximal actin disassembly is sufficient to drive this effect in vivo. Bottom insets show zoom-in on nascent myelin sheath at steps 1-3, with biomolecules (actin filaments, actin monomers, MBP) and plasma membrane drawn roughly to scale.

## DISUSSION

Compact myelin enables rapid impulse conduction by electrically insulating axons^2^. How oligodendrocytes form compact myelin—coordinating the requisite steps of cytoskeletal remodeling, membrane compaction, and wrapping—has remained unclear^13–15^. Here, by integrating tract-resolved developmental profiling, a live-cell compaction-mapping assay, and oligodendrocyte-targeted actin perturbations in vivo, we define a temporal and mechanistic relationship between actin filament disassembly and myelin growth. Our results support a working model in which sustained actin disassembly precedes active wrapping and promotes myelin wrapping/compaction by enabling MBP-associated membrane compaction (Fig. 7).

A central implication of our findings is that actin disassembly is unlikely to promote wrapping through repeated cycles of actin assembly and disassembly at the oligodendrocyte inner tongue, as proposed by actin cycling/ratcheting models^13,15,16^. In the thoracic gracile fasciculus, actin filament foci are frequent in nascent MBP+ sheath profiles early on, but decline sharply before the population transition to multilamellar compact myelin. We find little evidence for persistent actin filaments within sheaths during the phase of robust multilamellar growth. These timing data argue that actin filament loss is not simply a late endpoint of wrapping, but instead occurs upstream of it. In addition, our ultrastructural time course reveals that myelin growth is protracted over weeks to months, proceeding far more slowly than canonical actin-driven protrusion. Finally, experimentally biasing oligodendrocytes toward actin disassembly using DeAct genetic tools^43^ accelerates early myelin thickening in vivo rather than reducing it. Together, these observations favor a trigger-like transition in which actin filament disassembly precedes, and is permissive for, subsequent actin-independent mechanisms that support continued spiral growth.

Our data point to membrane compaction as a plausible downstream process coupled to actin loss. Compaction requires close apposition of the cytoplasmic leaflets and exclusion of cytoplasm and proteins from compact myelin^20,31^; thus, even sparse F-actin would be expected to sterically interfere with compaction given the 7 nm actin filament diameter^48,49^ relative to the 3 nm cytoplasmic gap of compact myelin^6,17,20^. Consistent with a compaction-permissive role for actin loss, prior work in cultured oligodendrocytes showed that pharmacologically promoting actin disassembly (e.g., cytochalasin D or latrunculin) increases the area of MBP-enriched membrane^14,15^, whereas disrupting endogenous actin-disassembly pathways (ADF/cofilin1 or gelsolin loss) increases noncompact myelin and decreases wrapping in vivo^14,15^. However, MBP enrichment alone is not a definitive readout of membrane compaction, as MBP is also present in non-compact regions such as the cell body and cytoplasmic channels^7,32^. We therefore developed a live-cell assay that maps MBP-associated compaction-like membrane states^21,40^ in cultured oligodendrocytes, enabling us to directly test how actin filaments relate to membrane compaction. Using correlative fluorescence imaging and bio-AFM on living cells, we show that EGFP-CAAX-excluded, membrane dye-positive domains correspond to nanoscale membrane thinning. These compaction zones are depleted of F-actin, and experimentally promoting actin disassembly increases their abundance and drives their expansion and fusion. In vivo, inducing actin filament disassembly in oligodendrocytes accelerates early myelin thickening, and this effect is strongest when actin disassembly is targeted to membrane-proximal regions, whereas perinuclear actin disassembly or increasing actin monomer levels does not phenocopy the phenotype. This spatial specificity supports the idea that the relevant actin population acts locally in myelin sheaths at or near apposing membranes, consistent with a compaction-permissive mechanism. Recent work has revealed that oligodendrocyte processes can form chains of myelin sheaths connected by thin cytoplasmic “paranodal bridges”—non-wrapping membrane segments that can persist into mature myelination^50^. The formation of these bridges, which contain F-actin but do not undergo wrapping, is consistent with our model in which local actin disassembly triggers compaction and wrapping: bridges may represent membrane domains where actin clearance is inhibited or delayed, rendering them refractory to MBP-mediated compaction.

How might compaction contribute to continued spiral growth? One possibility is that progressive membrane zippering^20^ and cytoplasmic extrusion during compaction alter the physical and geometrical constraints at the inner tongue^7^ in a way that facilitates continued advancement^14^. Several lines of evidence support tight coupling between compaction and wrapping: (1) non-compact myelin normally does not extend beyond ∼1–3 loose wraps around the axon, (2) myelin wrapping beyond the first few wraps does not initiate until after compaction begins^6,24^, (3) spiral growth of the inner tongue occurs simultaneously with compaction of outer layers^7^, and (4) in *Shiverer* mice that lack MBP and cannot compact myelin, wrapping fails to progress beyond a few loose turns^17,18^. Consistent with these observations, once compact myelin is evident in our developmental ultrastructural series, non-compact myelin rarely extends beyond approximately one turn around the axon, suggesting that wrapping and compaction remain closely linked as the sheath grows^6^. In this view, actin disassembly would function as an enabling step that permits MBP-associated compaction, which in turn supports continued wrapping/compaction as the sheath matures. Future experiments that directly track inner-tongue dynamics and compaction progression in vivo will be important for distinguishing whether compaction is merely permissive for wrapping or also contributes actively to the mechanics of spiral growth.

The relationship between MBP and actin disassembly may be more complex than a simple linear pathway. Our prior work demonstrated that MBP is itself required for actin disassembly in oligodendrocytes^14^: loss of MBP in *Shiverer* mice causes aberrant actin filament accumulation in ensheathing processes, and MBP competes with actin-disassembly factors such as cofilin-1 and gelsolin for binding to membrane PI(4,5)P2 in vitro, potentially releasing these factors from autoinhibition to enable local actin severing^14^. Together with the present findings that actin disassembly promotes MBP-associated compaction, these results suggest a feed-forward loop in which MBP binding to the membrane displaces and activates actin-disassembly proteins, local actin disassembly permits membrane compaction, and compaction in turn concentrates MBP to further drive the cycle. Such a mechanism could enable compaction zones to initiate at discrete sites and subsequently spread as a “traveling wave” across the sheath. This model is consistent with our live-cell observations that compaction zones expand and fuse over time (Fig. 3g,h) and with super-resolution imaging showing that MBP forms lattice-like domains bounded by actin filaments^14^. Testing this feed-forward model will require simultaneous tracking of MBP accumulation, actin disassembly, and membrane compaction at high spatial and temporal resolution, ideally in vivo during active myelin formation.

Beyond clarifying how actin remodeling is coupled to myelin growth, our compaction-mapping assay provides a platform to interrogate compaction dynamics directly. Notably, compaction zones in culture are highly dynamic, and we occasionally observe transient “decompaction-like” events in which previously compacted regions lose the compaction signature before stabilizing again. While this behavior remains to be validated in vivo and the underlying membrane and cytoplasmic changes require further definition, it suggests that compaction-like membrane states can be regulated on relatively short timescales. Such reversibility would be consistent with the idea that local antagonism of MBP-mediated adhesion—such as by proteins enriched in non-compact regions^32^—could tune the balance between compact and non-compact membrane states. Consistent with this idea, recent in vivo and ex vivo imaging studies indicate that myelin structure is more dynamic than previously appreciated, with activity modulating sheath swelling/decompaction-like pathology that can partially resolve over time^51^. Moreover, learning/plasticity-associated remodeling includes shortening of pre-existing myelin sheaths^52,53^, implying that mature sheaths can undergo rapid structural change that may require regulated transitions between compact and non-compact membrane states.

Our findings reveal a striking contrast with the mechanisms governing myelin sheath length. In our recent work, we demonstrated that actin filament assembly at sheath edges is necessary for longitudinal extension along axons: inducing actin disassembly reduced sheath length by ∼21% in vivo and blocked further elongation in myelinating cocultures^37^. Here, we show that the orthogonal growth axis—radial wrapping and thickening—is instead promoted by actin disassembly. This mechanistic divergence suggests that oligodendrocytes employ spatially segregated cytoskeletal programs to independently control myelin length versus thickness. This separation likely reflects distinct biophysical requirements: longitudinal extension resembles canonical cell migration requiring actin-based edge protrusion, whereas spiral wrapping/compaction involves progressive membrane zippering that is sterically incompatible with F-actin. Such modular control could enable activity-dependent tuning of myelin geometry^54^, with local calcium transients^55–58^ promoting actin assembly^59,60^ to tune myelin sheath lengths while other mechanisms (e.g., membrane remodeling^7,36,61^) separately tune sheath thickness/wrapping. Interestingly, actin disassembly has recently emerged as a pro-regenerative intervention in other CNS contexts^62,63^. Genetic or pharmacological promotion of actin depolymerization in retinal ganglion cells enhances axon regeneration after injury^63^, suggesting that modulating actin dynamics may represent a broader therapeutic strategy for CNS repair. In the context of remyelination, our findings suggest that promoting oligodendrocyte actin disassembly could accelerate myelin wrapping and compaction, whereas strategies that modulate actin assembly at sheath edges might preferentially enhance sheath length—implying that restoring these two growth modes may require distinct or combinatorial therapeutic approaches.

Several limitations should be considered when interpreting these results. First, our in vivo ultrastructural measurements quantify *g*-ratio and thickness across all axons within the analyzed tract rather than restricting analysis to transduced sheaths; thus, the magnitude of DeAct effects likely underestimates the cell-autonomous phenotype. Second, DeAct constructs provide a strong perturbation of actin dynamics^43^; determining how endogenous actin disassembly is regulated spatially and temporally within developing sheaths remains an important goal. Third, cultured oligodendrocyte sheets do not recapitulate all aspects of multilamellar sheaths around axons; compaction mapping therefore reports an MBP-associated compaction-like membrane state that is experimentally tractable, but necessarily reductionist.

In summary, our results support a model in which sustained actin filament disassembly precedes active wrapping and promotes myelin wrapping/compaction by enabling MBP-associated membrane compaction (Fig. 7). By providing both in vivo evidence for spatially constrained actin-dependent control of early myelin thickening and a live-cell platform to quantify compaction dynamics, this work also establishes an experimental system to define regulators of myelin compaction during development and to determine how compaction is perturbed in disease.

## ACKNOWLEDGMENTS

We thank current and past members of the Zuchero lab, Xiaowei Zhuang; and Julia Kaltschmidt, Kang Shen, Wah Chiu, and Aaron Gitler for discussion and support, and to Eva Carvalho for critical reading of the manuscript. We also thank the Stanford University Cell Sciences Imaging Core Facility for Transmission Electron Microscopy data collection, especially John Perrino and Ibanri Phanwar-Wood for their expertise in processing and staining EM samples (RRID:SCR_017787: supported by an ARRA Award Number 1S10RR026780-01 from the National Center for Research Resources. Its contents are solely the responsibility of the authors and do not necessarily represent the official views of the NCRR or the National Institutes of Health.). Electron microscopy image processing was supported in part by the grants: NN1 NSF 1707356 and NN2 NSF 2014862. We thank Nadia Makarova and Christina Newcomb at the Stanford University Cell Sciences Imaging Core Facility (RRID:SCR_017787) for training and technical assistance with atomic force microscopy. This work used the Bruker BioScope Resolve BioAFM, supported by NIH ORIP grant 1 S10 OD021514-01. We also thank Nicholas R. Wall and the Stanford Gene Vector and Virus Core (RRID:SCR_023250) for producing the AAVs used in this study. Images and diagrams were created using BioRender.com.

This project was supported by the Stanford Berry Postdoctoral Fellowship (H.K.); Stanford Wu Tsai Neurosciences Interdisciplinary Graduate Fellowship (K.W.); Stanford Medical Scientist Training Program (T32 GM007365-45; K.W. and M.H.C.); NINDS Diversity Supplements R01NS119823-01S1 and R01NS119823-04S1 (E.C.B., M.A.G., and J.B.Z.); Stanford Medicine Postbaccalaureate Experience in Research Program (E.C.B.); Stanford Bio-X Interdisciplinary Graduate Fellowship (M.H.C.); Wu Tsai Neurosciences Interdisciplinary Postdoctoral Scholarship (M.A.G.); Regina Casper Stanford Graduate Fellowship (M.I.); the National Multiple Sclerosis Society Harry Weaver Neuroscience Scholar Award (J.B.Z.); the McKnight Endowment Fund for Neuroscience (J.B.Z.); the Myra Reinhard Family Foundation (J.B.Z.); the Koret Family Foundation (J.B.Z.); and the National Institutes of Health R01NS119823 (J.B.Z.).

## AUTHOR CONTRIBUTIONS

J.B.Z. conceived the project and supervised the study. H.K., K.W., N.A., E.C.B., M.H.C., A.M., Y.M.S., M.A.G., M.I., D.M.J., J.B.Z. designed and performed experiments and analyzed data.

D.M.J. assisted with electron microscopy sample prep, imaging, and data analysis. H.K., K.W., N.A., M.H.C., M.A.G., M.I., and J.B.Z. generated key reagents. J.B.Z. obtained funding for the project. H.K., K.W., and J.B.Z. wrote the manuscript with input from all authors. All authors read and approved the final manuscript.

## METHODS

### Animals and ethics

All procedures involving animals were approved by Stanford University’s Institutional Animal Care and Use Committee (IACUC; also known as the Administrative Panel on Laboratory Animal Care, APLAC) and were conducted in accordance with National Institutes of Health guidelines. Stanford University’s laboratory animal care program is fully accredited by AAALAC International.

Mice were group-housed in a Stanford University animal facility under a 12:12 h light/dark cycle with ad libitum access to food and water, and were monitored by veterinary and animal care staff. Animals used in the study had not undergone prior procedures. C57BL/6J mice were used for all in vivo experiments unless otherwise noted and were obtained from Charles River (catalog #027). Previous studies have shown that myelination progresses more rapidly in male mice than in females. To minimize variability in developmental timing, we performed our time-course analysis of myelination between P4 and P90 in male animals. Both male and female mice were used in other experiments. For cell culture experiments, brains from both sexes were pooled to obtain sufficient cell numbers.

Mice were randomized for all procedures. First, the pilot experiment was conducted unblinded to make observations and determine the parameters for quantification. Subsequently, all microscopy and quantitative analyses were performed blinded to experimental condition and were only unblinded after analysis. Sample sizes (number of animals) are reported in the figure legends; no animals were excluded from analysis.

For tissue collection for immunohistochemistry, mice were anesthetized with ketamine (100 mg/kg) and xylazine (20 mg/kg) and transcardially perfused with phosphate-buffered saline (PBS) followed by 4% paraformaldehyde (PFA) (diluted in PBS from 16% PFA stock; Electron Microscopy Sciences). Alternatively, after anesthetizing with isoflurane, euthanasia was performed by decapitation. The spinal cords were then extracted from the spinal column by extrusion using a blunted needle, and tissues were immediately placed in the fixative after fluorescent examination. This allowed us to examine the spinal cords for construct expression after AAV injections, ensuring that the processed spinal cords were successfully injected and oligodendrocytes were expressing the target constructs. For electron microscopy experiments requiring chemical fixation, animals were perfused with PBS followed by Karlsson–Schultz fixative (KS fixative; 13 mM NaH2PO4, 87 mM Na2HPO4, 85.6 mM NaCl, 2.5% glutaraldehyde, and 4% PFA) and tissues were post-fixed in KS fixative at 4 °C prior to processing.

### AAV constructs, production, and neonatal intrathecal injections

#### AAV plasmid constructs

Adeno-associated virus (AAV) expression plasmids were generated in a pAAV backbone using the 1.9 kb MBP promoter (pMBP) to drive oligodendrocyte-lineage expression^35,36^. Constructs used in this study included: pMBP-EGFP-CAAX (control)^37^, pMBP-DeAct-GS1-P2A-EGFP-CAAX (“global/cytoplasmic” DeAct-GS1)^43^, pMBP-DeAct-GS1-EGFP-CAAX (membrane-tethered DeAct-GS1; no P2A)^37^, pMBP-EGFP-KASH (control), pMBP-DeAct-GS1-KASH (perinuclear-targeted^45,46^ DeAct-GS1), pMBP-Actin(R62D)-P2A-EGFP-CAAX (nonpolymerizable actin mutant^47^), and pCMV-Lifeact-mRuby3^41,42^. All constructs were created using InFusion cloning (Takara Bio) by annealing one or more DNA fragments into the parent plasmid. DNA fragments were produced using PCR or directly synthesized as mouse codon-optimized Infusion-competent “gene fragments without adaptors” (Twist Biosciences), in both cases with the 15 base pair overhangs required for InFusion reactions. All constructs were sequence-verified by full plasmid sequencing (Plasmidsaurus) both at the time of cloning and after midiprepping (Qiagen) for AAV production.

#### AAV production, storage, and titration

AAV-DJ serotype viruses were produced by the Stanford Neuroscience Gene Vector and Virus Core and stored at −80 °C until use. Viruses were thawed on ice immediately prior to injection and kept cold until use. Viral genomic titers were as follows:

| **Virus Name** | **Genomic titer (viral genomes/mL)** |
| --- | --- |
| AAV DJ pBZ-281 pMBP-EGFP-CAAX | 1.48E+13 |
| AAV DJ pBZ-283 pMBP-GS1-P2A-EGFP-CAAX | 2.75E+13 |
| AAV DJ pBZ-423 pMBP-GS1-EGFP-CAAX | 6.05E+12 |
| AAV DJ pBZ-364 pMBP-EGFP-KASH | 1.77E+13 |
| AAV DJ pBZ-368 pMBP-DeAct-GS1-KASH | 1.90E+13 |
| AAV DJ pBZ-342 pMBP-Actin(R62D)-P2A-EGFP-CAAX | 1.60E+13 |
| AAV DJ pBZ-235 pCMV-Lifeact-mRuby3 | 1.80E+13 |

#### Neonatal intrathecal injections

Neonatal (P0–P1) C57BL/6J mouse pups were cryoanesthetized on ice until unresponsive to toe pinch. AAVs were injected intrathecally into the lumbar spinal cord using a Hamilton syringe (Model 80308 701SN, Point Style 4, 32 G, 20 mm length, 12°). Virus was mixed with Trypan Blue to visualize successful delivery (1:2 with Trypan Blue), a volume of 2-3 µL was injected per pup. Successful injections were verified by observing a blue “stripe” indicating rostrocaudal spread of dye/virus within the spinal canal. Pups were recovered on a 34–37 °C warming surface until mobile and then returned to the home cage.

After euthanasia and recovery of spinal cords by extrusion, the spinal cords were inspected for construct expression by placing in a Mattek dish and visualization under a Zeiss Axio Observer Z1 inverted microscope, ensuring that the spinal cords were successfully injected. Spinal cords collected at the indicated ages were processed for downstream analyses (immunostaining, SiR-actin labeling, or electron microscopy) as described below.

### Electron microscopy

For the time course and AAV perturbation experiments (Figs. 1,2, 5, and 6), mice were perfused with PBS followed by Karlsson–Schultz fixative (KS fixative; composition in “Animals”). Thoracic spinal cords were dissected and post-fixed in KS fixative at 4 °C until EM processing (for at least 16-24 hrs). Alternatively, after anesthetizing with isoflurane, euthanasia by decapitation, the spinal cords were then carefully extruded from the spinal column with a blunted needle, and the tissues were promptly placed in the fixative. This process enabled us to verify the presence of construct expression in the spinal cords following AAV injections, confirming successful injection. Prior to electron microscopy (EM) preparation, thoracic spinal cord segments were identified as the region immediately posterior to the cervical enlargement. Approximately 1 cm of the thoracic spinal cord was dissected, and the posterior orientation was marked by an angled dorsoventral lesion. In experiments in which both histological staining and EM samples were obtained from the same spinal cord, the cord was transected at the T1 segment immediately after the cervical enlargement. The segment anterior to this cut was processed for transmission electron microscopy (TEM), whereas the posterior portion was processed for staining as described below. The most anterior thoracic segments were selected for analysis. Samples were processed for TEM in the Stanford Cell Sciences Imaging Facility as described in ^64^. Samples were washed twice in 0.1 M phosphate buffer (pH 7.3) at 4 °C and post-fixed in 2% osmium tetroxide prepared in the same buffer for 4 h at 4 °C. Samples were subsequently washed three times for 10 min each in 0.1 M phosphate buffer at 4 °C. Dehydration was done at 4 °C through a graded ethanol series consisting of 30%, 50%, 70%, and 95% ethanol (two 10-min incubations at each concentration), followed by four 10-min incubations in 100% ethanol. These steps were done on a rotator placed in a cold room at 4 °C. Samples were then transitioned to propylene oxide for 30 min at room temperature and infiltrated with propylene oxide: Epon (Embed 812 Hard) mixtures at ratios of 2:1 and 1:1 for 1 h each with rotation. Further infiltration was carried out in a 1:2 propylene oxide: Epon mixture overnight at room temperature, followed by infiltration in 100% Epon for 4 h at room temperature. Samples were embedded in labeled flat molds and allowed to equilibrate for 1 h prior to repositioning and polymerization. Polymerized blocks were trimmed, and semi-thin sections were prepared (200 nm) and stained with toluidine blue for light microscopic evaluation and to locate the dorsal column. Ultrathin sections were taken around 80nm using a UC7 (Leica, Wetzlar, Germany), picked up on formvar/carbon-coated 100 mesh Cu grids, stained for 40 seconds in 3.5% Uranyl Acetate in 50% Acetone, followed by staining in Sato’s Lead Citrate for 2 minutes. Sections were observed in the JEOL JEM-1400 120kV. Images were taken using a Gatan OneView 4k X 4k digital camera. Tiled image montages were obtained with SerialEM^65^, and images were opened with IMOD^66,67^.

#### EM classification and morphometry

Axons within the thoracic gracile fasciculus were categorized as (i) unmyelinated, (ii) ensheathed (defined as 1–3 loose, non-compact wraps), or (iii) wrapping/wrapped (defined as multilayered compact myelin containing major dense line-positive layers), as described in the Results. For *g*-ratio and thickness measurements, only regions/axons with well-preserved myelin ultrastructure (i.e., compact myelin without visible decompaction artifacts) were analyzed. Tissues exhibiting severe dehydration artifacts, including tearing and poorly defined axonal or myelin morphology, were excluded from analysis.

Axon diameter and myelin thickness were measured in Fiji/ImageJ by manual tracing or line measurements. Axon diameter was measured as the shortest distance between opposing axonal membranes that was perpendicular to the longest axis of the axon. This approach was used to account for oblique sectioning, as transverse sections that are not perfectly perpendicular to the horizontal plane of the spinal cord yield elliptical axonal profiles in TEM images. Myelin thickness was measured from the most intact, well-preserved, and compact region of the myelin sheath surrounding each axon. The *g-ratio* was calculated as the axon diameter divided by the axon diameter plus twice the myelin thickness. For each animal, measurements from individual axons were averaged to obtain a per-animal mean; statistical analyses were performed on per-animal means as indicated in the figure legends.

### Estimation of growth rate of myelin wrapping from EM time course

To quantify the rate of myelin wrapping during developmental myelination while minimizing bias from the continuous addition of newly initiated, thin sheaths, we analyzed the thickest 10% of myelin sheaths from the dorsal mouse spinal cord at each timepoint. This enrichment strategy increased the likelihood of sampling a relatively stable population of maturing sheaths across developmental stages. At each timepoint, we measured mean axon diameter and myelin thickness. We converted myelin thickness to the estimated number of wraps assuming a single-wrap thickness of 14.5 nm. We estimated total myelin spiral length using a flat-roll (Archimedean) spiral approximation, which is well-suited for compact myelin because successive wraps increase the sheath radius by a constant interval. We calculated spiral length (*L*) using the following formula:

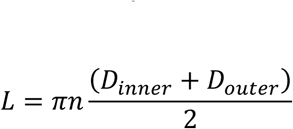

This simplifies to:

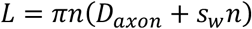

Where *n* = number of wraps, *D* = diameter, and *s_w_* = single wrap thickness.

Increases in spiral length can arise from two mechanisms: (i) addition of new wraps and (ii) radial expansion of existing wraps as axon caliber increases and new wraps displace older layers outward, increasing their circumference (spiraling leading edge/inner tongue akin to a runner circling a track of increasing diameter). We estimated the cumulative contribution of new wraps and radial expansion to spiral length increases at each timepoint interval using the following formulas:

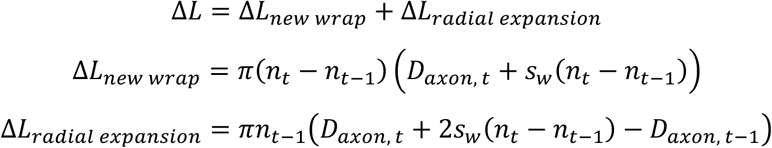

Although wraps added between timepoints also undergo radial expansion, it is difficult to isolate the contribution of radial expansion for these newly added wraps. We therefore attributed all length associated with new wraps to wrap addition. Accordingly, new-wrap values represent <u>upper bounds</u> on active wrapping, a conservative assignment given the expected minor contribution of radial expansion to newly deposited wraps. We plotted estimated total spiral length and the new-wrap component versus developmental time. Because total spiral length comprises new wraps plus radial expansion, the radial expansion component is represented by the difference between these curves. We fit each dataset with a logarithmic function, which appropriately models growth that decelerates over time. Time-resolved growth rates for total spiral length and the new-wrap component were computed by differentiating the respective fitted curves.

Finally, we compared the calculated rate of myelin wrapping to published rates of several forms of canonically actin-driven cell motility/process extension, including human neutrophils/lymphocytes undergoing fast amoeboid motility in culture (∼600 μm/hr^68–70)^, cancer (glioma) cells (∼50 μm/hr^71^), neuronal growth cones (∼25 μm/hr^72^), fibroblasts during wound healing (∼8 μm/hr^73^). Rate numbers were derived from: https://bionumbers.hms.harvard.edu/search.aspx

### Primary oligodendrocyte precursor cell (OPC) purification and oligodendrocyte culture

Primary oligodendrocyte precursor cells (OPCs) were purified by immunopanning from postnatal rodent brains as previously described^14,34^. Briefly, OPCs were isolated from P5–P7 Sprague-Dawley rat pups, seeded onto poly-D-lysine (PDL)-coated tissue culture dishes, and allowed to recover and proliferate before experiments.

All plasticware was coated with 0.01 mg/mL PDL (Sigma P6407) in water. Glass coverslips were coated with 0.01 mg/mL PDL prepared in 150 mM boric acid pH 8.4 (“PDL-borate”) and diluted to 1× in water prior to coating.

To proliferate OPCs, cells were maintained in serum-free defined “DMEM-SATO” base medium supplemented with 4.2 µg/mL forskolin (Sigma F6886), 10 ng/mL PDGF (Peprotech 100-13A), 10 ng/mL CNTF (Peprotech 450-02), and 1 ng/mL neurotrophin-3 (NT-3; Peprotech 450-03) at 37 °C and 10% CO_2_. To induce oligodendrocyte differentiation, cultures were switched to DMEM-SATO base medium containing 4.2 µg/mL forskolin, 10 ng/mL CNTF, and 40 ng/mL triiodothyronine (T3; Sigma T6397). Unless otherwise noted, “DIV” refers to days in differentiation medium (days after switching from OPC proliferation medium to differentiation medium).

For compaction-mapping and actin experiments, OPCs were plated at 5500 cells/well (8-well dishes, Ibidi 80807) or 8500 cells/dish (35 mm dish with 14 mm glass coverslip, MatTek P35GC-1.5-14-C.S) and transduced at the time of plating with AAVs expressing fluorescent reporters and/or DeAct constructs under the pMBP promoter (see “AAV constructs, production, and neonatal intrathecal injections”). Media was partially exchanged every 2–3 days with half-media changes. Oligodendrocytes were analyzed at the indicated days in differentiation medium (e.g., DIV3 for live compaction mapping and DIV7 for fixed-cell assays), as specified in the figure legends.

#### Viral transduction and expression timing in culture

For oligodendrocyte culture experiments, AAVs were added at the time of OPC plating into differentiation medium at a dilution of 1:6000 (empirically determined to be lowest dilution that leads to expression in >80% of cells with no cytotoxicity; see *AAV production, storage, and titration* above for exact titers). For experiments comparing DeAct-GS1 versus control, cells were transduced with either pMBP-DeAct-GS1-P2A-EGFP-CAAX or the matched pMBP-EGFP-CAAX control. For Lifeact experiments, oligodendrocytes were transduced with pCMV-Lifeact-mRuby3.

#### Pharmacological manipulations in culture

For acute actin disassembly experiments, latrunculin A (LatA; Invitrogen L12379) was prepared as a 1 mM stock solution in DMSO (stored in small aliquots with desiccant at −20 °C for no longer than 6 months) and diluted to 2x (500 nM) in imaging solution prior to use. Oligodendrocytes were imaged at 1 frame/hr, with DMSO-only controls processed in parallel. LatA was added immediately before the second frame to a final concentration of 250 nM.

#### Fixation and immunostaining of cultured oligodendrocytes

At the specified day in differentiation medium, cells were fixed in 4% PFA in PBS for 15 min at room temperature and washed 3× with PBS. Cells were permeabilized with 0.1% Triton X-100 in PBS for 3 min. F-actin was stained with Alexa Fluor 568-conjugated phalloidin (Thermo Scientific A12380) diluted 1:143 in PBS for 15 min. Coverslips were mounted in Fluoromount G with DAPI (SouthernBioTech, 0100-20).

### Compaction mapping and live-cell imaging in cultured oligodendrocytes

#### Compaction mapping (EGFP-CAAX exclusion with membrane dye continuity)

To visualize MBP-associated membrane compaction in cultured oligodendrocytes, cells were transduced at plating with an AAV expressing EGFP-CAAX under the pMBP promoter. To label the plasma membrane independent of protein content, cultures were stained with MemGlow 640 (Cytoskeleton, Inc., #MG04) at a final concentration of 20 nM.

For fixed-cell experiments, MemGlow was applied in PBS after fixation at room temperature for at least 20 min prior to imaging. Because MemGlow signal is lost upon permeabilization, Memglow and EGFP-CAAX fluorescence images were acquired prior to permeabilization and aligned to post-permeabilization images using custom Python scripts employing Fourier-based rigid registration.

For live imaging, oligodendrocytes were incubated in pre-warmed imaging medium containing MemGlow for at least 20 min at 37 °C and imaged without washout.

Compaction zones were operationally defined as membrane regions that were (i) MemGlow-positive (continuous membrane) and (ii) depleted of EGFP-CAAX relative to surrounding membrane. Oligodendrocytes expressing EGFP-CAAX and labeled with MemGlow are referred to as “compaction-mapped oligodendrocytes.”

#### Live imaging of compaction dynamics

Live imaging was performed on a Zeiss Axio Observer Z1 inverted microscope equipped with a Zeiss Axiocam 506 monochrome 6-megapixel camera, a stage top incubator (Okolab, H301-K-Frame) set to 37°C, and a digital gas blender (Okolab, CO2-UNIT-3L) set to 10% CO_2_ during image acquisition. Prior to imaging, culture medium was exchanged through three partial media swaps for differentiation medium prepared with FluoroBrite DMEM (Thermo Fisher A1896701) in place of standard DMEM to reduce background fluorescence and phototoxicity (hereafter: “imaging medium”). Cells were imaged using a 20x/0.8 NA objective. Single-plane time-lapse images of EGFP-CAAX and MemGlow were acquired every hour for ∼10 hours using Zen Blue software, using gentle illumination settings to minimize phototoxicity and bleaching.

For time-lapse datasets, photobleaching correction was performed using histogram matching in Fiji/ImageJ prior to segmentation. Cell outlines were defined from MemGlow signal, and compact versus non-compact membrane regions were classified based on EGFP-CAAX depletion within the cell outline, as described in “Image analysis.”

#### Live imaging of actin dynamics relative to compaction

To monitor actin dynamics during compaction, oligodendrocytes were transduced with pCMV-Lifeact-mRuby3. Lifeact and compaction-mapping channels were imaged at the same interval as above (every hour). For analyses of actin intensity at compaction onset, regions of interest were defined in membrane areas that transitioned from non-compact to compact during the time series, and Lifeact intensity was quantified before and after compaction onset as described in “Image analysis.”

#### Correlative fluorescence imaging and bio-atomic force microscopy (bio-AFM)

##### Correlative imaging workflow

Correlative fluorescence imaging and atomic force microscopy (AFM) were performed on fixed compaction-mapped oligodendrocytes to measure membrane height in regions classified as compact versus non-compact by fluorescence. Oligodendrocytes expressing pMBP-EGFP-CAAX and labeled with MemGlow were imaged first by fluorescence microscopy to identify EGFP-CAAX-excluded, MemGlow-positive compaction zones. Cells were then immediately fixed and the same cells and regions of interest were relocated for AFM using the gridded dish reference system (MatTek P35G-1.5-14-C-GRD) and the AFM’s integrated brightfield microscope. AFM imaging was performed within hours of fixation.

##### AFM instrumentation and acquisition

AFM measurements were acquired using a Bruker BioScope Resolve in PBS at room temperature. A soft cantilever suitable for live-cell measurements (PEAKFORCE-HIRS-F-A cantilever (Bruker; spring constant k = 0.4 N/m, resonant frequency 165 kHz, tip radius ∼1 nm) was used. Cantilever spring constants were calibrated using the thermal tune method prior to imaging (nominal k = 0.4 N/m).

Topography scans were collected in PeakForce QNM mode at a resolution of 256×256, scan size of 20-500 μm and scan rate of 0.25-0.5 Hz using a PeakForce setpoint of 0.5-2 nN.

##### Height measurement and definition of compact regions

Membrane height was measured from AFM topography and processed using Mountains SPIP (Digital Surf, Besancon, France). Height data were flattened using line-by-line leveling with a first-order polynomial fit, scan lines affected by significant artifacts were replaced by adjacent line interpolation, and height values were zeroed to the mean height of the glass substrate. AFM images were overlaid with their corresponding immunofluorescence images using up to 10 manually selected registration points. Line profiles (3 per biological sample) were extracted, and height and fluorescence data were exported for quantification.

Using a custom Python script, line profiles were baseline-corrected to the mean height of the glass substrate within each line. A histogram was generated from the distribution of this height data. Individual pixels were then classified by height: 12–16 nm as compact and >17 nm as non-compact. EGFP-CAAX intensity was quantified and compared between compact and non-compact pixels.

Each biological replicate consisted of an independent oligodendrocyte culture preparation. For each replicate, 1-2 cells were scanned and 3 line profiles per cell were analyzed, as specified in the figure legends.

### Actin labeling and imaging in tissue

#### Phalloidin and MBP immunostaining of spinal cord sections (developmental time course)

For the developmental actin time course, thoracic spinal cords were collected at the indicated postnatal ages and processed for immunofluorescence. Animals were anesthetized and transcardially perfused with PBS followed by 4% PFA (see “Animals”). Spinal cords were post-fixed in 4% PFA for overnight. For a second cohort of animals at all indicated ages (P4-P24), after isoflurane anesthesia and decapitation, spinal cords were extruded out of the spinal column using a blunted needle and hydraulic pressure of PBS, and tissues were immediately placed in 4% PFA in PBS for 45 mins. Tissues were washed in PBS, cryoprotected in 30% sucrose at 4 °C 3-7 days, embedded in OCT, and sectioned on a cryostat (Leica CM3050S) at 10 µm. Sections were mounted on Superfrost Plus slides, dried, and stored at −80 °C until staining.

Slides were dried at room temperature for 15-30 mins. Sections were permeabilized and blocked in 0.1% Triton X-100, 10% goat serum in PBS for 1 h, and incubated with primary antibodies (including rat anti-MBP) diluted in 1:100, 1% donkey serum in PBST overnight at 4 °C. After PBS washes, sections were incubated with Alexa Fluor-conjugated secondary antibodies for 2 h at room temperature. F-actin was labeled with Alexa Fluor-conjugated phalloidin (Invitrogen Molecular Probes Alexa Fluor 647 Phalloidin A22287) at 1:50 for 2 hrs. Slides were mounted in Molecular Probes ProLong Gold Antifade Mountant, P10144, with DAPI and imaged as described in “Microscopy and image acquisition.”

#### Confocal imaging

Confocal imaging was performed on a Zeiss LSM 880 using Zeiss Zen Blue software. For high-resolution imaging and quantitative actin measurements, samples were imaged using 63×/1.4 NA oil objective with Airyscan confocal microscopy. Laser power and acquisition settings were held constant within each experiment, and imaging was performed blinded to condition.

#### Quantification of actin foci in MBP+ sheath profiles

MBP+ sheath profiles were identified as ring-like MBP immunofluorescence structures in spinal cord cross-sections. Actin foci were scored as discrete phalloidin-positive puncta within MBP-defined sheath regions using a consistent intensity threshold per experiment. Data were aggregated to per-animal means prior to statistical testing.

### STORM imaging of F-actin relative to myelin sheath markers

STORM was used to localize F-actin relative to myelin sheath markers in thoracic spinal cord tissue at the indicated ages. Except as noted, coverslip preparation, buffers, instrumentation, imaging acquisition, image processing and visualization were then performed as described in ^74^. Code developed to process and analyze STORM reconstructions is available at https://github.com/SpeerLab/STORM_analysis.

First, following tissue perfusion, tissue was post-fixed in 3% Paraformaldehyde/0.1% Glutaraldehyde overnight at 4C. Tissue was then washed in PBS, and sectioned using a vibratome at 120μm thickness. Samples were collected using the Tokuyasu method and were first cryoprotected in 2.3M Sucrose overnight at 4C. 400nm thick sections were cut on an ultra-cryomicrotome (UC7 Ultramicrotome with FC7 cryo-attachment, Leica Microsystems, using an ultra Jumbo diamond knife, Diatome). Samples the then picked up in a 1:1 mixture of 2.3M sucrose and 2% methylcellulose, and placed in a coated (0.5% gelatin/0.05% chromium potassium sulfate) 8-well chambered coverslip for staining and imaging.

Images were collected in Storm imaging buffer (10% glucose/17.5 μM glucose oxidase/708 nM catalase/10 mM MEA/10 mM NaCl/200 mM Tris) using a custom built microscope and oblique incident angle illumination as described. For each image reconstruction, 50,000 frames (60Hz) were collected at each wavelength and centroids were localized across a 100μm x 100μm field of view. STORM data were acquired to achieve ∼10–20 nm lateral (x–y) localization precision and reconstructed images were rendered at 15 nm per pixel. Conventional diffraction-limited reference images were acquired for each section at ∼158 nm per pixel and used for registration and thresholding-based filtering of STORM localizations. For registration, two-color widefield conventional images were collected at the same time and chromatically corrected using a fiducial based (715/755 FluoSpheres - Life Technologies) polynomial mapping. Each super-resolution image was then drift corrected using a normalized cross-correlation and then rigidly aligned to the conventional image.

All STORM processing and analysis were performed blinded to age/condition. Image contrast was adjusted linearly for visualization within each dataset and channel; therefore, fluorescence intensities were not compared quantitatively across samples.

### SiR-actin target engagement assay in AAV-injected spinal cords

#### SiR-actin labeling in spinal cord sections

For labeling actin filaments in vivo in AAV-injected animals, actin puncta within EGFP-labeled myelin sheath profiles were visualized using the cell-permeable and fluorogenic SiR-actin probe (Cytoskeleton, CY-SC001). After isoflurane anesthesia and euthanasia with decapitation, spinal cords were squirted out of the spinal column using a blunted needle, and tissues were immediately placed in 4% PFA for a brief 45-minute fixation. Thoracic spinal cords were collected, and to preserve EGFP fluorescence, they were processed without sucrose cryoprotection and without thin cryosectioning, then sectioned at 120 µm using a vibratome. A brief 1-hour blocking was done with 10% goat serum. Tissues were then incubated with SiR-actin (10 µM) and Hoechst (Invitrogen H3570, 1:1000) containing PBS overnight at 4 °C on a cold room rotator. Z-stacks spanning the full thickness of the EGFP+ myelin rings were obtained with Airyscan confocal microscopy in the entire region of the dorsal thoracic gracile fasciculus. SiR-actin puncta overlapping EGFP+ sheaths were quantified blinded to condition and summarized as the percentage of EGFP+ sheath profiles containing one or more distinct SiR-actin puncta. Each dot in plots represents the per-animal quantification, and analyses were performed blinded to condition.

### In vivo AAV expression/penetrance and construct localization controls

#### Quantification of in vivo myelin labeling penetrance

To assess in vivo labeling penetrance of pMBP-driven EGFP reporters, thoracic spinal cords were collected at the indicated ages from animals injected at P0–P1 with pMBP-EGFP-CAAX or pMBP-DeAct-GS1-P2A-EGFP-CAAX. Spinal cords were fixed in 4% PFA, cryoprotected in 30% sucrose, embedded in OCT, and cryosectioned at 10 µM. Because EGFP fluorescence is quenched in PFA-perfused, sucrose-cryoprotected tissue, a delipidation step was performed to enable GFP detection by antibody staining. Slides were soaked in 100% ethanol for 10 min, followed by three 10-min washes in PBS, prior to the blocking step. Sections were immunostained for MBP (Abcam ab7349; knockout-validated^14^; 1:100) and GFP (Santa Cruz, #sc-9996, 1:500) and imaged by confocal microscopy with identical acquisition settings for each age group.

MBP+ myelin rings were identified in spinal cord cross-sections and scored for EGFP positivity. Labeling penetrance was calculated as the fraction of MBP+ myelin rings that were EGFP+ for each animal.

### Construct localization controls in cultured oligodendrocytes (KASH targeting^45^)

To validate perinuclear targeting of KASH-tagged constructs, primary oligodendrocytes were transduced with pMBP-EGFP-KASH or pMBP-DeAct-GS1-KASH and fixed at DIV4. Cells were stained with phalloidin and CellMask Blue to delineate the soma/perinuclear compartment and imaged by widefield epifluorescence (Zeiss Axio Observer Z1 inverted microscope, 20x objective).

Perinuclear enrichment of EGFP signal was quantified as the ratio of mean EGFP intensity within a soma/perinuclear ROI (defined using CellMask Blue) to mean EGFP intensity across the whole cell.

### Image analysis and quantification

For in vivo experiments, the first pilot experiment was conducted unblinded to make observations and determine the parameters for quantification. Subsequently, all image acquisition and quantitative analyses were performed blinded to experimental condition and unblinded only after analysis. Image processing was performed in Fiji/ImageJ and Python unless otherwise noted.

#### Image preprocessing

For datasets requiring alignment (including fixed-cell multichannel images and time-lapse sequences), images were registered using Fourier-based rigid registration implemented in custom Python scripts. For time-lapse datasets, EGFP-CAAX and MemGlow channels were corrected for photobleaching in Fiji using the Bleach Correction plugin (Histogram Matching). Unless otherwise stated, analyses were performed on single optical sections.

#### Definition of regions of interest (ROIs)

- *Cultured oligodendrocytes (compaction mapping)*: Individual cells were selected by manually drawing a ROI around each oligodendrocyte in Fiji. Cell outlines were defined from MemGlow signal. Background regions were defined from areas lacking cells within the same field of view.
- *MBP “ring” ROIs (spinal cord sections)*: MBP+ sheath profiles were identified as ring-like MBP immunofluorescence structures in spinal cord cross-sections. In Supplementary Fig 3, MBP ring ROIs were generated by ilastik^75^. For each section, ROIs were extracted without viewing the phalloidin/SiR-actin channel to minimize bias.
- *EGFP+ sheath profiles (SiR-actin target engagement)*: EGFP+ sheath profiles were identified in dorsal column white matter in thick sections. Sheath profiles were scored in 3D z-stacks by first background-subtracting from the actin channel using the mean intensity of approximately 500 of the darkest pixels in each image. The GFP and F-actin channels were then merged, and the brightness/contrast of the actin channel was adjusted uniformly across all images using fixed display values (2027/2050). Myelin sheaths were quantified by measuring the presence of F-actin puncta associated with individual myelin sheaths.

In Supplementary Fig. 3C, the overlap between ilastik-derived MBP ring masks and phalloidin staining was quantified without experimenter or thresholding bias using a custom ImageJ macro. Multichannel Airyscan images were imported into ImageJ and split into individual fluorescence channels, and the phalloidin (F-actin) channel was selected for analysis.

Background subtraction was performed by estimating background intensity from approximately 500 of the darkest pixels in each image and subtracting this value from the analysis channel to account for slide-to-slide variability in nonspecific staining intensity.

Binary masks of MBP rings were generated in ilastik using a trained classifier and imported into ImageJ. Images were then analyzed using a systematic intensity threshold sweep applied to the actin channel, ranging from 0 to 3,000 in increments of 50. For each threshold, a binary actin mask was created, and overlap with the MBP ring mask was quantified using the ImageJ “AND” function. All analyses were fully automated and batch-processed using identical parameters for all images; automation was routinely validated by manually auditing analysis.

#### Segmentation of compact and non-compact membrane in culture

Compaction zones were operationally defined as MemGlow-positive membrane regions depleted of EGFP-CAAX relative to surrounding membrane.

- *Fixed-cell segmentation*: Fixed-cell images were segmented using an Otsu-based thresholding approach applied to EGFP-CAAX intensity within the MemGlow-defined cell outline. To improve segmentation robustness, images were pre-processed by Gaussian filtering and by clipping pixel values across a range of outlier thresholds prior to applying Otsu thresholding. The final segmentation for each cell was selected by visual comparison to the raw images.
- *Time-lapse segmentation*. Time-lapse images were segmented using Labkit (Fiji) for pixel classification. Classifiers were trained using line annotations across compact and non-compact regions and applied across timepoints. Segmentations were manually reviewed and corrected as needed to ensure temporal consistency.

For time-lapse datasets, “newly compacted” membrane between frames was computed as the set difference between compact masks in consecutive timepoints. The number of compaction zones was defined as the number of connected components in the compact mask (8-connected), and mean compaction zone area was computed as total compact area divided by zone number.

#### Quantification of actin signal in culture

Phalloidin and Lifeact channels were background corrected by subtracting the median pixel intensity of each image (which corresponded to background). For fixed-cell phalloidin analyses, mean phalloidin intensity was quantified within compact versus non-compact membrane masks for each cell. For DeAct-GS1 validation in culture, mean phalloidin intensity per cell was measured within the MemGlow-defined cell outline.

For Lifeact time-lapse experiments, simple-ratio correction was applied to correct for increasing whole-cell Lifeact expression over time. To account for cell-to-cell variability in Lifeact expression, Lifeact intensity values were normalized by each cell’s mean across all timepoints. Lifeact intensities within compact and non-compact regions pixels were quantified over time. To assess actin dynamics leading up to “at compaction onset,” pixels that compacted during the time-lapse were identified and Lifeact intensity was quantified over the 5 h preceding and including the compaction event. Trajectories were aligned per pixel so compaction events were registered at t = 0, and compared to trajectories of never-compacting pixels within 50 px sharing the same t = 0.

#### Quantification of in vivo myelin ultrastructure

Axon diameter, myelin thickness, and *g*-ratio were measured from EM images in Fiji/ImageJ. For each animal, measurements from individual axons were averaged to obtain a per-animal mean; statistical tests were performed on per-animal means as described in “Statistics.” SuperPlots^76^ depict individual axons and per-animal means.

#### Software

Analyses were performed using Fiji/ImageJ (version 2.14.0/1.54f)^77^, Labkit (version 0.3.11), Python (version 3.10.9; key packages: bioio 3.2.0, scikit-image 0.19.3, NumPy 1.24.2, SciPy 1.10.1, pandas 1.5.3, matplotlib 3.5.3, seaborn 0.12.2, statsmodels 0.13.5), and GraphPad Prism (version 11.0.0). Custom scripts used for analysis are available upon request.

### Statistics

All experiments were randomized and analyzed blinded to experimental condition, and datasets were unblinded only after completion of quantification. Data are presented as mean ± SEM unless otherwise indicated. The unit of replication (biological replicate) and exact sample sizes are reported in the figure legends and/or Supplementary Data 1.

For imaging-based cell culture experiments, biological replicates corresponded to independent cell preparations. For in vivo experiments, biological replicates corresponded to individual animals. Where measurements were obtained from many individual structures within an animal (e.g., axons), values were first averaged within each animal to generate a per-animal mean; statistical tests were performed on per-animal means. These distributions are visualized using SuperPlots^76^, in which individual observations are shown alongside per-replicate means.

Normality was assessed using Shapiro-Wilk tests and visual inspection of Q-Q plots. Two-group comparisons were performed using paired or unpaired two-tailed t tests as appropriate. For comparisons involving repeated measurements over time (e.g., compaction dynamics in time-lapse imaging), two-way repeated-measures ANOVA was used with per-timepoint post hoc pairwise tests and multiple-comparison correction using the Benjamini-Hochberg false discovery rate. Exact statistical tests and P values for each analysis are reported in the figure legends and/or source data.

Statistical analyses were performed using GraphPad Prism (version 11.0.0) and Python (Scipy and statsmodels). A significance threshold of α = 0.05 was used unless otherwise noted.

### Data, reagent, and code availability

Source data supporting the findings of this study are provided with the paper. Custom analysis code, including scripts for image segmentation and quantification, will be made available at https://github.com/wukathryn/OL_compaction_analysis upon publication. DNA constructs generated for this study will be deposited to Addgene at the time of publication. Additional materials and information are available from the corresponding author upon reasonable request.

## REFERENCES

1. Almeida, R. G. & Lyons, D. A. On Myelinated Axon Plasticity and Neuronal Circuit Formation and Function. J. Neurosci. Off. J. Soc. Neurosci. 37, 10023–10034 (2017).

2. Stadelmann, C., Timmler, S., Barrantes-Freer, A. & Simons, M. Myelin in the Central Nervous System: Structure, Function, and Pathology. Physiol. Rev. 99, 1381–1431 (2019).

3. Xin, W. & Chan, J. R. Myelin plasticity: sculpting circuits in learning and memory. Nat. Rev. Neurosci. 16, 668–13 (2020).

4. Peters, A. FURTHER OBSERVATIONS ON THE STRUCTURE OF MYELIN SHEATHS IN THE CENTRAL NERVOUS SYSTEM. 20, 281–296 (1964).

5. Bunge, M. B., Bunge, R. P. & Pappas, G. D. Electron microscopic demonstration of connections between glia and myelin sheaths in the developing mammalian central nervous system. http://jcb.rupress.org/content/12/2/448/tab-metrics (1962).

6. Hildebrand, C., Remahl, S., Persson, H. & Bjartmar, C. Myelinated nerve fibres in the CNS. Prog. Neurobiol. 40, 319–384 (1993).

7. Snaidero, N. et al. Myelin membrane wrapping of CNS axons by PI(3,4,5)P3-dependent polarized growth at the inner tongue. Cell 156, 277–290 (2014).

8. Snaidero, N. & Simons, M. The logistics of myelin biogenesis in the central nervous system. Glia 65, 1021–1031 (2017).

9. Simons, M. & Nave, K.-A. Oligodendrocytes: Myelination and Axonal Support. Cold Spring Harb. Perspect. Biol. 8, a020479 (2015).

10. Chang, K.-J., Redmond, S. A. & Chan, J. R. Remodeling myelination: implications for mechanisms of neural plasticity. Nat. Neurosci. 19, 190–197 (2016).

11. Hartline, D. K. & Colman, D. R. Rapid conduction and the evolution of giant axons and myelinated fibers. Curr. Biol. CB 17, R29–35 (2007).

12. Bauer, N. G., Richter-Landsberg, C. & Ffrench-Constant, C. Role of the oligodendroglial cytoskeleton in differentiation and myelination. Glia 57, 1691–1705 (2009).

13. Brown, T. L. & Macklin, W. B. The Actin Cytoskeleton in Myelinating Cells. Neurochem. Res. 6, 22741–10 (2019).

14. Zuchero, J. B. et al. CNS myelin wrapping is driven by actin disassembly. Dev. Cell 34, 152–167 (2015).

15. Nawaz, S. et al. Actin filament turnover drives leading edge growth during myelin sheath formation in the central nervous system. Dev. Cell 34, 139–151 (2015).

16. Samanta, J. & Salzer, J. L. Myelination: Actin Disassembly Leads the Way. Dev. Cell 34, 129–130 (2015).

17. Rosenbluth, J. Central myelin in the mouse mutant shiverer. J. Comp. Neurol. 194, 639–648 (1980).

18. Shine, H. D., Readhead, C., Popko, B., Hood, L. & Sidman, R. L. Morphometric analysis of normal, mutant, and transgenic CNS: correlation of myelin basic protein expression to myelinogenesis. J. Neurochem. 58, 342–349 (1992).

19. Aggarwal, S., Yurlova, L. & Simons, M. Central nervous system myelin: structure, synthesis and assembly. Trends Cell Biol. 21, 585–593 (2011).

20. Bakhti, M., Aggarwal, S. & Simons, M. Myelin architecture: zippering membranes tightly together. Cell. Mol. Life Sci. CMLS 71, 1265–1277 (2014).

21. Aggarwal, S. et al. Myelin membrane assembly is driven by a phase transition of myelin basic proteins into a cohesive protein meshwork. PLoS Biol. 11, e1001577 (2013).

22. Matthews, M. A. & Duncan, D. A quantitative study of morphological changes accompanying the initiation and progress of myelin production in the dorsal funiculus of the rat spinal cord. J. Comp. Neurol. 142, 1–22 (1971).

23. Foran, D. R. & Peterson, A. C. Myelin acquisition in the central nervous system of the mouse revealed by an MBP-Lac Z transgene. J. Neurosci. Off. J. Soc. Neurosci. 12, 4890–4897 (1992).

24. Bunge, R. P. Glial cells and the central myelin sheath. Physiol. Rev. 48, 197–251 (1968).

25. Watkins, T. A., Emery, B., Mulinyawe, S. & Barres, B. A. Distinct stages of myelination regulated by gamma-secretase and astrocytes in a rapidly myelinating CNS coculture system. Neuron 60, 555–569 (2008).

26. Czopka, T., Ffrench-Constant, C. & Lyons, D. A. Individual Oligodendrocytes Have Only a Few Hours in which to Generate New Myelin Sheaths In-Vivo. Dev. Cell 25, 599–609 (2013).

27. Zeller, N. K., Behar, T. N., Dubois-Dalcq, M. E. & Lazzarini, R. A. The timely expression of myelin basic protein gene in cultured rat brain oligodendrocytes is independent of continuous neuronal influences. J. Neurosci. Off. J. Soc. Neurosci. 5, 2955–2962 (1985).

28. Xu, K., Zhong, G. & Zhuang, X. Actin, spectrin, and associated proteins form a periodic cytoskeletal structure in axons. Science 339, 452–456 (2013).

29. He, J. et al. Prevalent presence of periodic actin-spectrin-based membrane skeleton in a broad range of neuronal cell types and animal species. Proc. Natl. Acad. Sci. U. S. A. 113, 6029–6034 (2016).

30. Huang, B., Babcock, H. & Zhuang, X. Breaking the diffraction barrier: super-resolution imaging of cells. 143, 1047–1058 (2010).

31. Aggarwal, S. et al. A Size Barrier Limits Protein Diffusion at the Cell Surface to Generate Lipid-Rich Myelin-Membrane Sheets. Dev. Cell 21, 445–456 (2011).

32. Snaidero, N. et al. Antagonistic Functions of MBP and CNP Establish Cytosolic Channels in CNS Myelin. Cell Rep. 18, 314–323 (2017).

33. Beniac, D. R. et al. Three-dimensional structure of myelin basic protein. I. Reconstruction via angular reconstitution of randomly oriented single particles. J. Biol. Chem. 272, 4261–4268 (1997).

34. Dugas, J. C. & Emery, B. Purification of oligodendrocyte precursor cells from rat cortices by immunopanning. Cold Spring Harb. Protoc. 2013, (2013).

35. Gow, A., Friedrich, V. L. & Lazzarini, R. A. Myelin basic protein gene contains separate enhancers for oligodendrocyte and Schwann cell expression. J Cell Biol 119, 605–616 (1992).

36. Lam, M. et al. CNS myelination requires VAMP2/3-mediated membrane expansion in oligodendrocytes. Nat. Commun. 13, 5583 (2022).

37. Iyer, M. et al. Oligodendrocyte calcium signaling promotes actin-dependent myelin sheath extension. Nat. Commun. 15, 265 (2024).

38. Collot, M. et al. MemBright: A Family of Fluorescent Membrane Probes for Advanced Cellular Imaging and Neuroscience. Cell Chem. Biol. 26, 600–614.e7 (2019).

39. Collot, M., Boutant, E., Lehmann, M. & Klymchenko, A. S. BODIPY with Tuned Amphiphilicity as a Fluorogenic Plasma Membrane Probe. Bioconjug. Chem. 30, 192–199 (2019).

40. Kattnig, D. R., Bund, T., Boggs, J. M., Harauz, G. & Hinderberger, D. Lateral self-assembly of 18.5-kDa myelin basic protein (MBP) charge component-C1 on membranes. Biochim. Biophys. Acta 1818, 2636–2647 (2012).

41. Riedl, J. et al. Lifeact: a versatile marker to visualize F-actin. Nat. Methods 5, 605–607 (2008).

42. Bajar, B. T. et al. Improving brightness and photostability of green and red fluorescent proteins for live cell imaging and FRET reporting. Sci. Rep. 6, 20889–12 (2016).

43. Harterink, M. et al. DeActs: genetically encoded tools for perturbing the actin cytoskeleton in single cells. Nat. Methods 14, 479–482 (2017).

44. Lukinavičius, G. et al. Fluorogenic probes for live-cell imaging of the cytoskeleton. Nat. Methods 11, 731–733 (2014).

45. Swiech, L. et al. In vivo interrogation of gene function in the mammalian brain using CRISPR-Cas9. Nat. Biotechnol. 33, 102–106 (2015).

46. Ostlund, C. et al. Dynamics and molecular interactions of linker of nucleoskeleton and cytoskeleton (LINC) complex proteins. J. Cell Sci. 122, 4099–4108 (2009).

47. Posern, G., Sotiropoulos, A. & Treisman, R. Mutant actins demonstrate a role for unpolymerized actin in control of transcription by serum response factor. Mol. Biol. Cell 13, 4167–4178 (2002).

48. Hanson, J. & Lowy, J. The structure of F-actin and of actin filaments isolated from muscle. J. Mol. Biol. 6, 46-IN5 (1963).

49. von der Ecken, J., et al. Structure of the F-actin-tropomyosin complex. Nature 519, 114–117 (2015).

50. Call, C. L. et al. Flexible ensheathment of axons enables myelination of complex CNS networks. Nature 10.1038/s41586-026-10312-1 (2026) doi:10.1038/s41586-026-10312-1.

51. Arafa, D. et al. Myelin sheaths in the central nervous system can withstand damage and dynamically remodel. Science 391, eadr4661 (2026).

52. Yang, S. M., Michel, K., Jokhi, V., Nedivi, E. & Arlotta, P. Neuron class–specific responses govern adaptive myelin remodeling in the neocortex. Science 370, (2020).

53. Bacmeister, C. M. et al. Motor learning drives dynamic patterns of intermittent myelination on learning-activated axons. Nat. Neurosci. 25, 1300–1313 (2022).

54. Osso, L. A. & Hughes, E. G. Dynamics of mature myelin. Nat. Neurosci. 27, 1449–1461 (2024).

55. Braaker, P. N. et al. Activity-driven myelin sheath growth is mediated by mGluR5. Nat. Neurosci. 28, 1213–1225 (2025).

56. Dereddi, R. R. et al. Oligodendrocyte mechanotransduction channel TMEM63A regulates myelin sheath geometry. Neuron 114, 699–723.e11 (2026).

57. Halford, J. et al. TMEM63A, associated with hypomyelinating leukodystrophies, is an evolutionarily conserved regulator of myelination. Proc. Natl. Acad. Sci. U. S. A. 122, e2507354122 (2025).

58. Young, A. R. et al. CNS Myelin Sheath Lengths Locally Scale to Axon Diameter via Piezo1. BioRxiv Prepr. Serv. Biol. 2025.11.18.689045 (2025) doi:10.1101/2025.11.18.689045.

59. Brown, T. L., Hashimoto, H., Finseth, L. T., Wood, T. L. & Macklin, W. B. PAK1 Positively Regulates Oligodendrocyte Morphology and Myelination. J. Neurosci. 41, 1864–1877 (2021).

60. Leisner, T. M., Liu, M., Jaffer, Z. M., Chernoff, J. & Parise, L. V. Essential role of CIB1 in regulating PAK1 activation and cell migration. J. Cell Biol. 170, 465–476 (2005).

61. Wu, J. et al. Nonvesicular lipid transfer drives myelin growth in the central nervous system. Nat. Commun. 15, 9756 (2024).

62. Tedeschi, A. et al. ADF/Cofilin-Mediated Actin Turnover Promotes Axon Regeneration in the Adult CNS. Neuron 103, 1073–1085.e6 (2019).

63. Li, L. et al. Actin depolymerization promotes axon regeneration by restoring axonal mitochondrial transport in mouse models of optic neuropathy. Sci. Transl. Med. 18, eadw0908 (2026).

64. Mobius, W. et al. Electron microscopy of the mouse central nervous system. Methods Cell Biol. 96, 475–512 (2010).

65. Mastronarde, D. N. Automated electron microscope tomography using robust prediction of specimen movements. J. Struct. Biol. 152, 36–51 (2005).

66. Mastronarde, D. N. Dual-axis tomography: an approach with alignment methods that preserve resolution. J. Struct. Biol. 120, 343–352 (1997).

67. Kremer, J. R., Mastronarde, D. N. & McIntosh, J. R. Computer visualization of three-dimensional image data using IMOD. J. Struct. Biol. 116, 71–76 (1996).

68. Bear, J. E. & Haugh, J. M. Directed migration of mesenchymal cells: where signaling and the cytoskeleton meet. Curr. Opin. Cell Biol. 30, 74–82 (2014).

69. Fowell, D. J. & Kim, M. The spatio-temporal control of effector T cell migration. Nat. Rev. Immunol. 21, 582–596 (2021).

70. Gruler, H. New insights into directed cell migration: characteristics and mechanisms. Nouv. Rev. Fr. Hematol. 37, 255–265 (1995).

71. Prag, S. et al. NCAM regulates cell motility. J. Cell Sci. 115, 283–292 (2002).

72. Harris, W. A., Holt, C. E., Smith, T. A. & Gallenson, N. Growth cones of developing retinal cells in vivo, on culture surfaces, and in collagen matrices. J. Neurosci. Res. 13, 101–122 (1985).

73. Jonkman, J. E. N. et al. An introduction to the wound healing assay using live-cell microscopy. Cell Adhes. Migr. 8, 440–451 (2014).

74. Sigal, Y. M., Speer, C. M., Babcock, H. P. & Zhuang, X. Mapping Synaptic Input Fields of Neurons with Super-Resolution Imaging. 163, 493–505 (2015).

75. Berg, S. et al. ilastik: interactive machine learning for (bio)image analysis. Nat. Methods 16, 1226–1232 (2019).

76. Lord, S. J., Velle, K. B., Mullins, R. D. & Fritz-Laylin, L. K. SuperPlots: Communicating reproducibility and variability in cell biology. J. Cell Biol. 219, (2020).

77. Schindelin, J., et al. Fiji: an open-source platform for biological-image analysis. Nat. Methods 9, 676–682 (2012).

