## Supplementary Materials for "Actin disassembly triggers CNS myelin compaction and wrapping"

Contents include the following:

- Supplemental Figures and Figure Legends for Supplementary Figs. 1–6.
- Legends for Supplementary Videos 1-4.

### SUPPLEMENTARY FIGURES AND LEGENDS

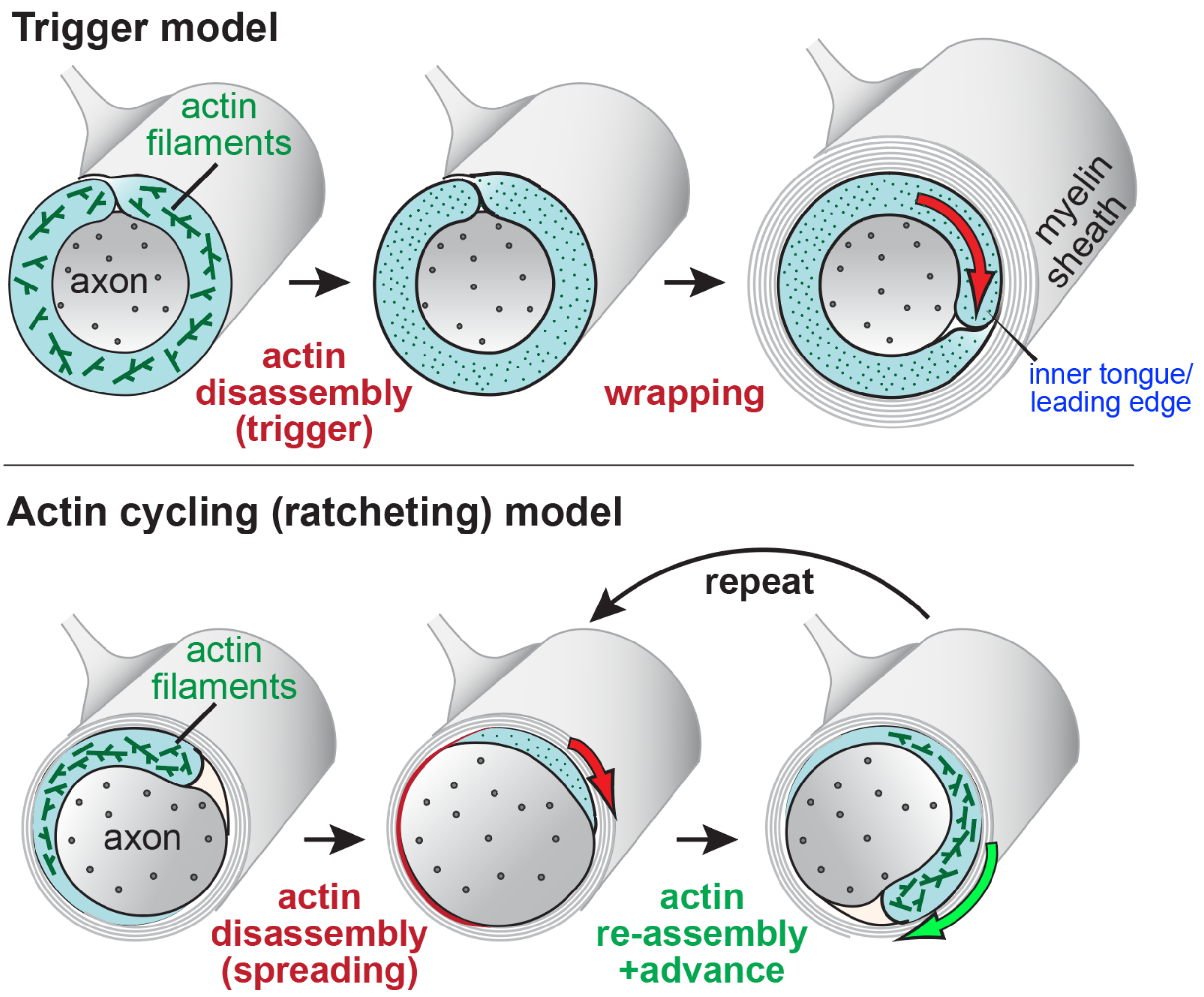


**Supplementary Fig. 1 | Competing models for how actin remodeling could advance myelin wrapping (related to Fig. 1).**

**Top**, Trigger model. During ensheathment, F-actin is present within the oligodendrocyte process. A developmental actin disassembly event acts as a trigger, after which wrapping proceeds without requiring sustained actin assembly at the leading edge.

**Bottom**, Actin cycling (ratcheting) model. Wrapping is driven by repeated cycles of actin disassembly and re-assembly at the inner tongue/leading edge that incrementally advance the oligodendrocyte membrane around the axon.

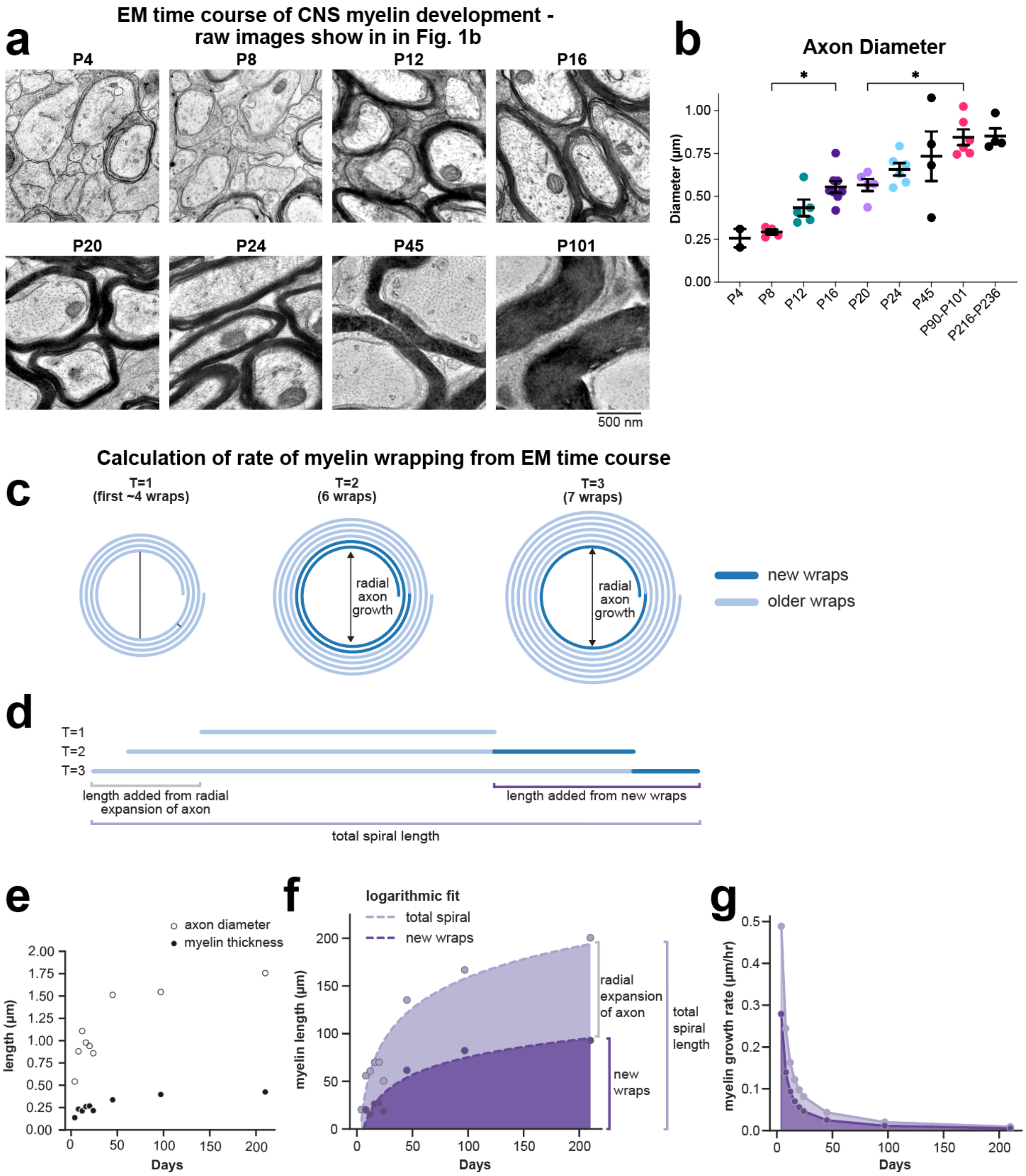


**Supplementary Fig. 2 | Additional characterization of thoracic gracile fasciculus development by EM (related to Fig. 1).**

**a**, Unprocessed representative EM images corresponding to the raw micrographs used for the false-colored examples shown in Fig. 1b. Scale bar, 500 nm.

**b**, Average axon diameter across development measured from gracile fasciculus EM cross-sections. Data are shown as mean ± SEM with individual animals plotted as points. Statistical tests and exact P values are reported in the Methods/source data. Sample sizes (animals) are provided in Supplementary Data 1.

**c-g**, Calculations of myelin wrapping growth rate show that developmental myelin wrapping is a slow, protracted process.

**c**, Conceptual schematic illustrating myelin spiral growth across three developmental timepoints (T=1, 2, 3). New wraps (dark blue) are added progressively to the inner surface while pre-existing wraps (light blue) undergo radial expansion as the axon grows and inner wraps displace older layers outward.

**d**, Proportional representation of the above myelin spirals unraveled at each timepoint, showing relative contributions of radial expansion (light blue) and new wraps (dark blue) to total spiral length increases.

**e**, Developmental trajectories of mean axon diameter (open circles) and mean myelin thickness (filled circles) used to estimate wrap number and spiral length, based on our EM time course of developmental myelination in the thoracic gracile fasciculus (Fig. 1).

**f**, Total myelin spiral length (light purple) and contributions from new wraps (dark purple markers), with corresponding logarithmic fits (dashed lines). Radial expansion is represented by the difference between these curves.

**g**, Instantaneous growth rate of myelin wrapping derived from the derivative of logarithmic fits.

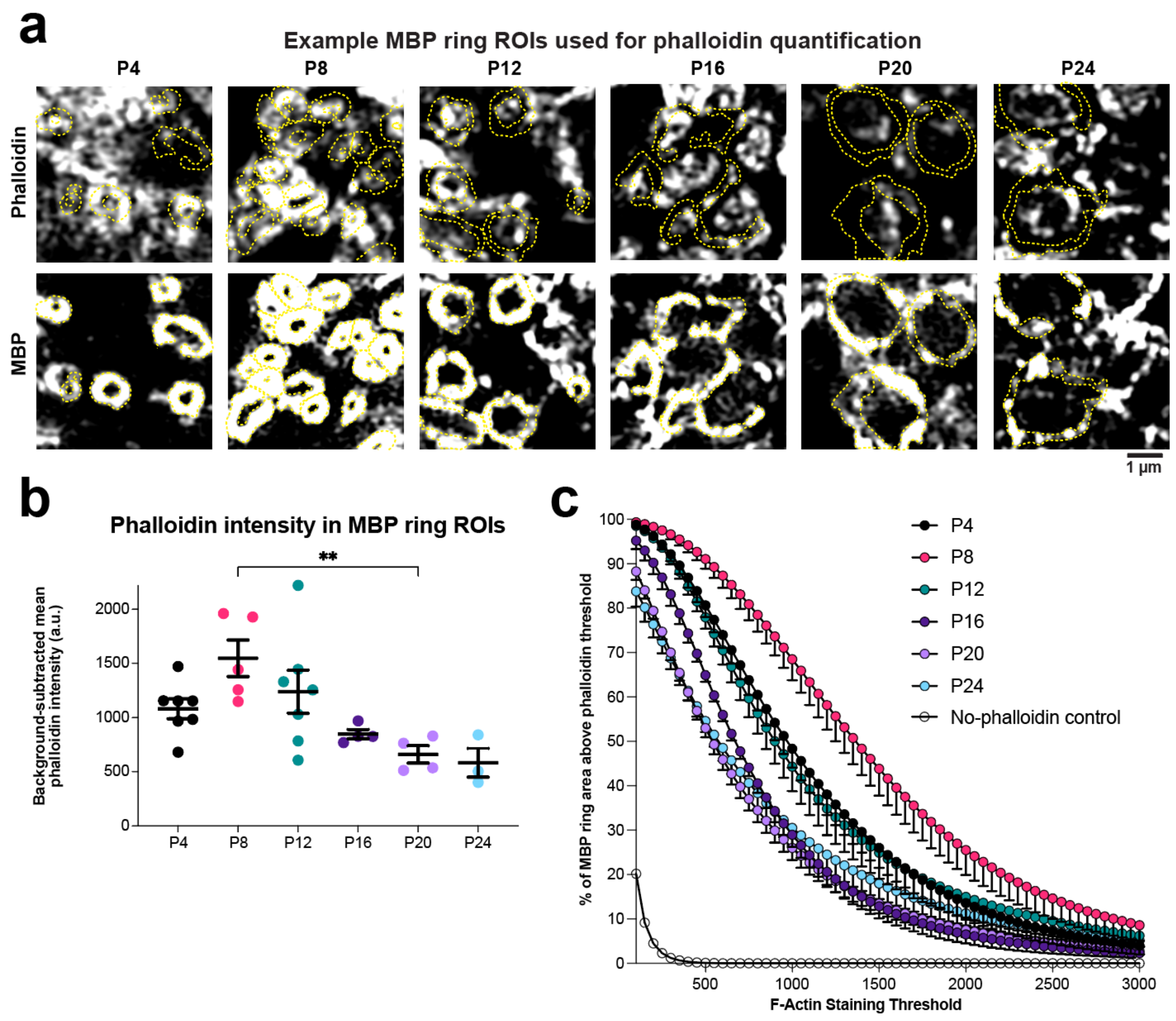


**Supplementary Fig. 3 | Quantification strategy and robustness for measuring F-actin signal in MBP ring ROIs (related to Fig. 2).**

**a**, Representative examples of MBP "ring" ROIs definition used for phalloidin quantification across postnatal ages. Yellow outlines indicate MBP ring ROIs. Scale bars, 1 µm.

**b**, Developmental time course of background-subtracted phalloidin (F-actin) intensity within MBP ring ROIs. Phalloidin intensity was measured within MBP ring ROIs following background subtraction (see Methods), then averaged within each animal to generate one value per animal (each dot = animal; mean ± SEM shown; n = 3–7 animals per age). Statistics: one-way ANOVA with Tukey test (brackets/asterisks as shown).

**c**, Threshold-sensitivity analysis for phalloidin signal within MBP ring ROIs. Phalloidin images were thresholded across a range of values and the percentage of MBP ring area above the phalloidin threshold was calculated for each animal. Curves show mean ± SEM across animals for each age. Open circles indicate a no-phalloidin control to account for tissue autofluorescence.

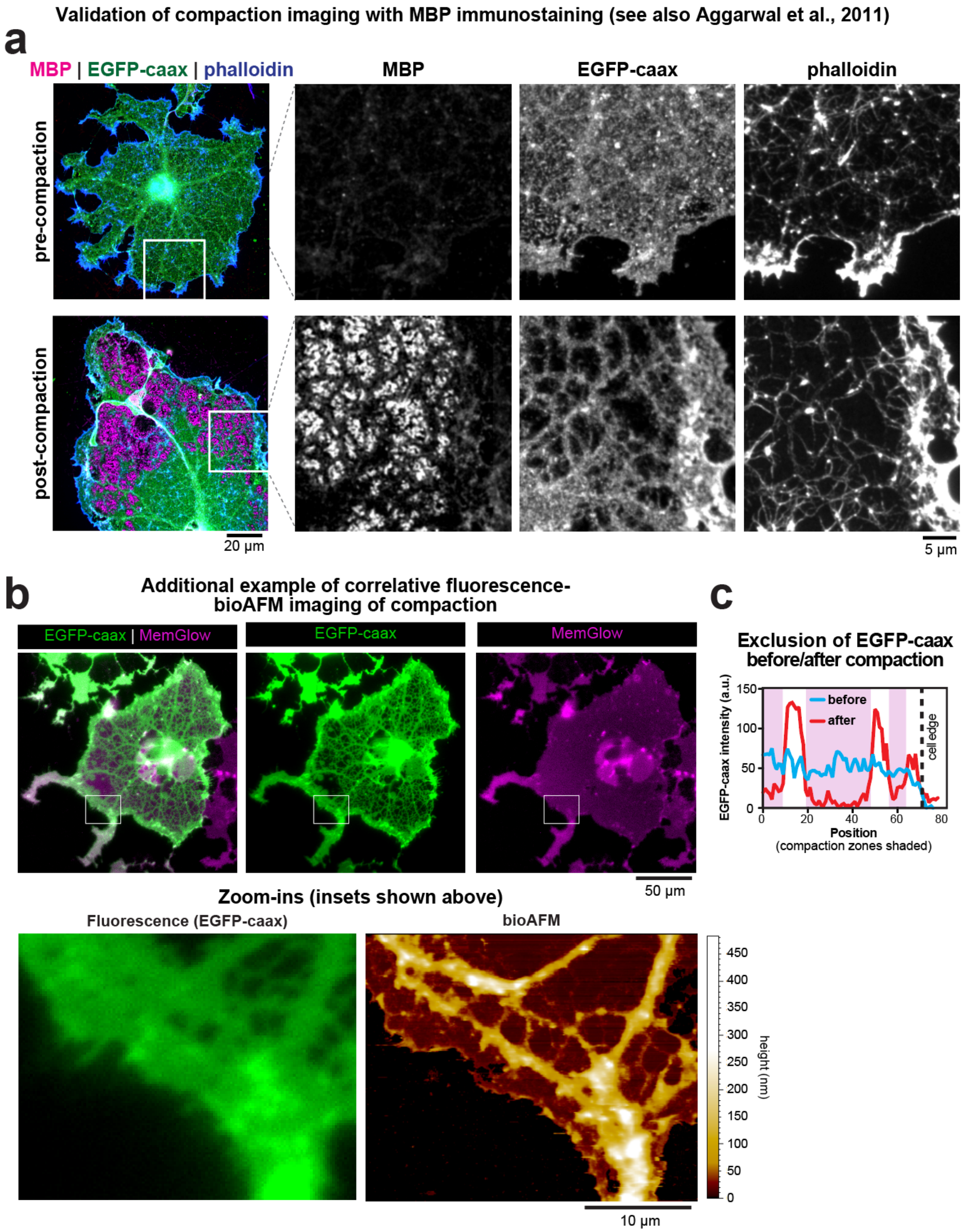


**Supplementary Fig. 4 | Validation of compaction imaging with MBP immunostaining and additional correlative bio-AFM example (related to Fig. 3).**

**a**, Representative 4-day-differentiated OLs before (“pre-compaction”) and after (“post-compaction”) formation of EGFP-CAAX-excluded membrane domains, stained for MBP and phalloidin. Insets show that MBP is enriched in EGFP-CAAX-excluded regions, as shown previously[31]. Scale bars: left, 20 μm; right, 5 μm.

**b**, Additional example of correlative fluorescence imaging (EGFP-CAAX, MemGlow) and bio-AFM height mapping of compaction zones. Box indicates the region shown at higher magnification (insets of paired fluorescence and bio-AFM shown at bottom). Height scale shown at right. Scale bars, 50 μm for top full-cell micrographs; 10 μm for bottom zoomed-in insets.

**c**, Line scan of movie stills before and after compaction of a representative oligodendrocyte shows exclusion of EGFP-CAAX from compaction zones and its enrichment in non-compact regions of the oligodendrocyte.

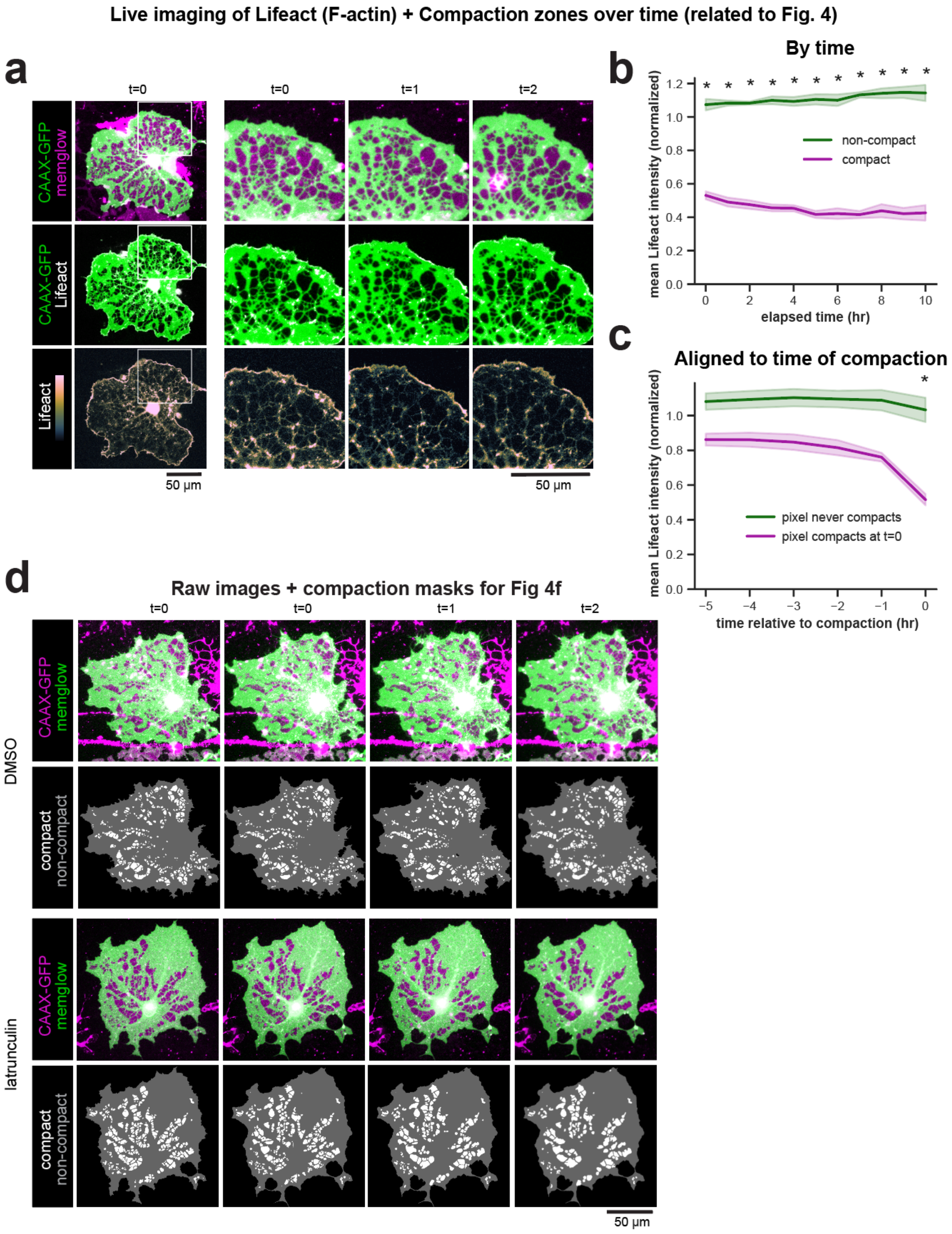


**Supplementary Fig. 5 | Live actin imaging and raw images underlying compaction masks (related to Fig. 4).**

**a**, Live imaging of a compaction-mapped OL expressing Lifeact-mRuby3 (to label actin filaments; hereafter “Lifeact”) showing lower actin signal in compact regions compared with non-compact regions over time. Plot to right shows mean actin intensity (a.u.) in compact versus non-compact regions (mean ± SEM; significance as indicated) over time. Scale bar, 50 μm.

**b**, Actin intensity is consistently lower in compact versus non-compact regions of cells. Mean Lifeact intensity in compact (purple) versus non-compact (green) pixels of each cell, plotted across elapsed time. Lifeact intensity values are normalized to each cell's mean Lifeact intensity across all timepoints. Lines and bands show mean ± SEM; N = 3 independent biological replicates. Two-way repeated-measures ANOVA; main effect of time, $p = 0.73$ (n.s.); main effect of group, $p = 3.7 \times 10^{-4}$; time × group interaction, $p = 0.0013$. Per-timepoint paired two-tailed t-tests, Benjamini–Hochberg FDR-corrected: significant at all tested timepoints t = 0–10 h ($p_{FDR}$ = 0.011, 0.0042, 0.0034, 0.0014, 0.0034, 0.0014, $9.1 \times 10^{-4}$, $9.1 \times 10^{-4}$, $9.1 \times 10^{-4}$, 0.0013, 0.0037).

**c**, Local actin signal drops around time of compaction. Mean Lifeact intensity leading up to compaction in pixels destined to compact (purple) versus neighboring non-compacting control pixels (green). Compaction events occurring at different absolute frames are time-aligned such that t = 0 marks the time of compaction for each pixel. Control pixels are time-matched, non-compacting pixels within 50 px of a compacting pixel. Lifeact intensity values are normalized to each cell's mean Lifeact intensity across all timepoints. Lines and bands show mean ± SEM; N = 3 independent biological replicates. Two-way repeated-measures ANOVA; main effect of time, $p = 1.5 \times 10^{-4}$; main effect of group, $p = 0.054$ (n.s.); time × group interaction, $p = 4.5 \times 10^{-6}$. Per-timepoint paired two-tailed t-tests, Benjamini–Hochberg FDR-corrected: significant at t = 0 h ($p_{FDR} = 0.037$); not significant at t = −5 to −1 h ($p_{FDR} = 0.10$).

**d**, Raw EGFP-CAAX/MemGlow fluorescence images and corresponding compaction masks from the latrunculin (LatA) time-course experiment (DMSO control versus latrunculin). Scale bar, 50 μm.

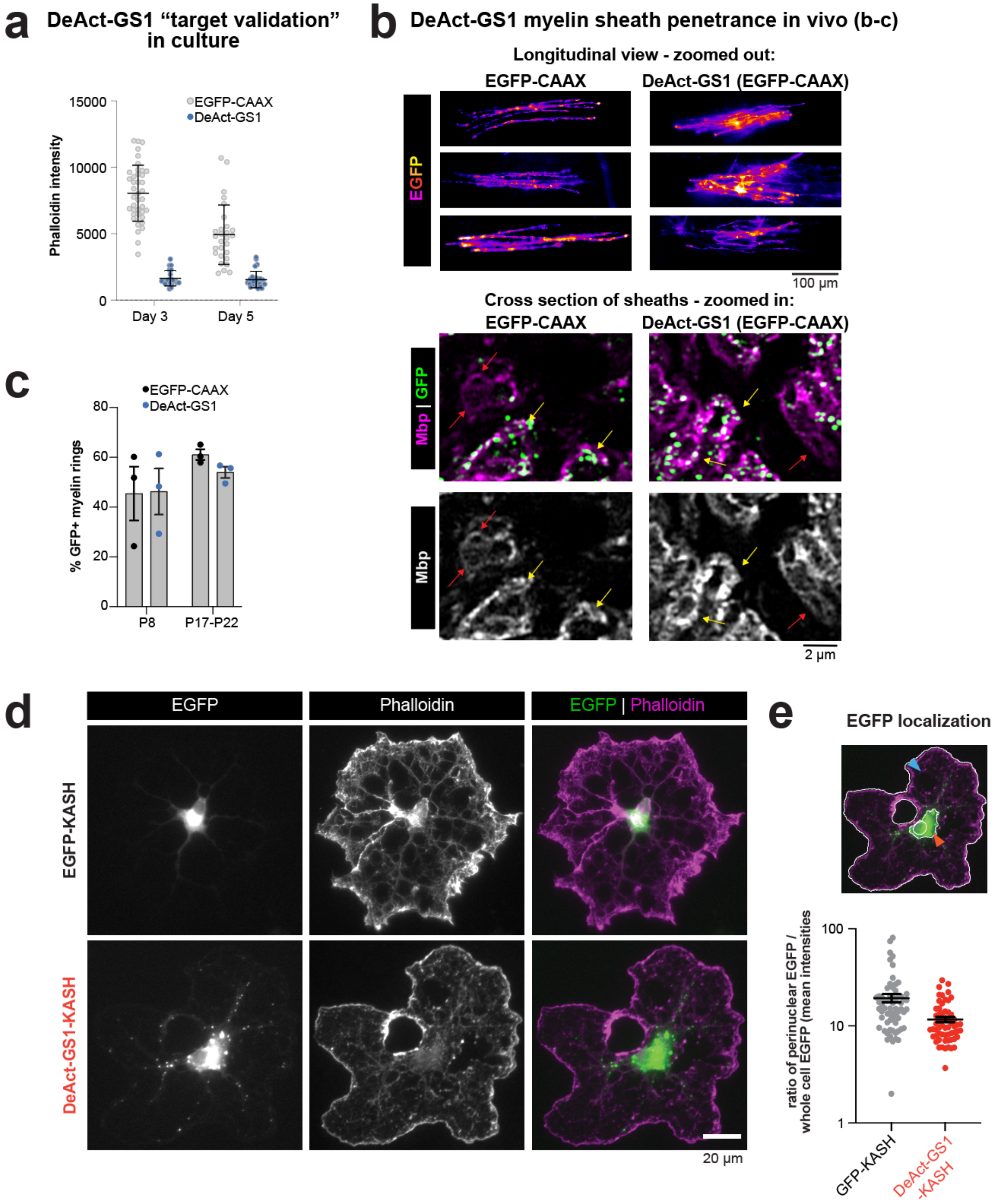


**Supplementary Fig. 6 | Validation and controls for in vivo DeAct perturbations (related to Figs. 5–6)**

**a**, DeAct-GS1 reduces F-actin in cultured oligodendrocytes. Primary oligodendrocytes expressing EGFP-CAAX (control; gray) or DeAct-GS1 (magenta) were fixed at 3- and 5-day-differentiated and stained with phalloidin. Plot shows mean phalloidin intensity per cell (each dot

= cell; N = 1 biological replicate). (See also prior culture validation of DeAct-GS1 in Iyer et al. 2022 Supplementary Figure 6, and Harterink et al. 2017 Figure 1)

**b**, Representative in vivo expression and myelin labeling. Maximum-intensity projections (top) of EGFP signal in thoracic dorsal column from animals injected with pMBP-EGFP-CAAX (left) or pMBP-GS1-P2A-EGFP-CAAX (“DeAct-GS1 (EGFP-CAAX)”, right). Bottom, representative thoracic spinal cord cross-sections delipidated[72] and immunostained for MBP (magenta) with EGFP (green) showing examples of MBP+ myelin rings that are EGFP+ (yellow arrows) or EGFP− (red arrows).

**c**, Myelin sheath labeling penetrance in vivo. Quantification of the percentage of MBP+ myelin rings that were EGFP+ at P8 and P17–P22 for pMBP-EGFP-CAAX and pMBP-GS1-P2A-EGFP-CAAX AAVs. Each dot represents the mean from one animal (N = 3 animals per condition/time point); bars indicate mean ± SEM.

**d**, KASH localization control in cultured oligodendrocytes. Representative 4-day-differentiated oligodendrocytes expressing EGFP-KASH (top) or DeAct-GS1-KASH (bottom) stained with phalloidin. Left, EGFP; middle, phalloidin; right, merge.

**e**, Quantification of perinuclear enrichment for KASH constructs. Perinuclear EGFP localization was quantified as the ratio of mean EGFP intensity in a soma/perinuclear ROI (defined by CellMaskBlue staining) to mean EGFP intensity across the whole cell. Each dot = cell; bars indicate mean ± SEM. Y-axis is log-scaled.

Scale bars: b, 100 μm (top) and 2 μm (bottom); d, 20 μm.

### LEGENDS FOR SUPPLEMENTARY VIDEOS

**Supplementary Video 1. | Live-imaging and compaction mapping of cultured oligodendrocytes (related to Fig. 3)**

Video shows live imaging of a representative 3-day-differentiated compaction-mapped OL, imaged every 1 hr for 10 hrs. Left is fluorescence with EGFP-CAAX (green) and MemGlow (magenta). Right shows segmentation of compact (white) and non-compact (gray) membrane regions by EGFP-CAAX exclusion. Scale bar, 25 µm.

**Supplementary Video 2. | Dynamics of compaction zones during their formation (related to Fig. 3)**

Video shows live imaging of another representative 3-day-differentiated compaction-mapped OL, imaged every 1 hr for 11 hrs. Left panel shows fluorescence with EGFP-CAAX (green) and MemGlow (magenta). Right panel shows segmentation of compact (white) and non-compact (gray) membrane regions by EGFP-CAAX exclusion. Compaction zones that will decompact in the next frame (orange) and newly appearing compaction zones (blue) are highlighted. Upper panel shows the whole OL (scale bar, 25 µm); lower panel shows a zoomed-in view of the region outlined by the white box (scale bar, 5 µm).

**Supplementary Video 3. | Two-color live-imaging of compaction and actin dynamics (related to Fig. 4c)**

Video shows live imaging of the same cell as in Supplementary Video 1, with co-expressed Lifeact, imaged every 1 hr for 10 hrs. Left shows fluorescence with EGFP-CAAX (green) and Lifeact (blue); right shows Lifeact alone. Scale bar, 25 µm.

**Supplementary Video 4. | Dynamics of compaction zones following acute actin disassembly with latrunculin (related to Fig. 4f)**

Video shows live imaging of a representative 3-day-differentiated compaction-mapped OL, imaged every 1 hr for 12 hrs. Movie starts at -1 hr (1 h prior to latrunculin addition); latrunculin was added at t = 0 h (note minor, transient decompaction at time of addition, which also occurred with DMSO addition). Left is fluorescence with EGFP-CAAX (green) and MemGlow (magenta). Right shows segmentation of compact (white) and non-compact (gray) membrane regions by EGFP-CAAX exclusion. Scale bar, 25 µm.
